# Low-dimensional genotype-fitness mapping across divergent environments suggests a limiting functions model of fitness

**DOI:** 10.1101/2025.04.05.647371

**Authors:** Olivia M. Ghosh, Grant Kinsler, Benjamin H. Good, Dmitri A. Petrov

## Abstract

A central goal in evolutionary biology is to be able to predict the effect of a genetic mutation on fitness. This is a major challenge because fitness depends both on phenotypic changes due to the mutation, and how these phenotypes map onto fitness in a particular environment. Genotype, phenotype, and environment spaces are extremely large and complex, rendering bottom-up prediction difficult. Here we show, using a large collection of adaptive yeast mutants, that fitness across a set of lab environments can be well-captured by low-dimensional linear models of abstract genotype-phenotype-fitness maps. We find that these maps are low-dimensional not only in the environment where the adaptive mutants evolved, but also in divergent environments. We further find that the genotype-phenotype-fitness spaces implied by these maps overlap only partially across environments. We argue that these patterns are consistent with a “limiting functions” model of fitness, whereby only a small number of limiting functions can be modified to affect fitness in any given environment. The pleiotropic side-effects on non-limiting functions are effectively hidden from natural selection locally, but can be revealed globally. These results combine to emphasize the importance of environmental context in genotype-phenotype-fitness mapping, and have implications for the predictability and trajectory of evolution in complex environments.

---

A fundamental challenge in evolutionary biology is developing a predictive framework that links genetic variation to phenotypic traits and, ultimately, to evolutionary fitness (3, 12, 14, 24, 44, 45, 50). One major obstacle is the sheer size of genotype space (9, 29, 35). Each genetic variant can modify molecular phenotypes, which in turn affect higher-level phenotypes, eventually causing macroscopic changes to organisms. Building bottom-up, causal chains across these phenotypic levels is extremely difficult given the vast network of interactions at every level of biological organization. Of particular interest in evolutionary biology is understanding how genetic changes affect phenotypes that matter to fitness (17, 37). Furthermore, the environment can modulate how genotypes map onto phenotypes, and their mapping onto fitness (4, 10, 43). Hence, a genotype-phenotype-fitness map must contend with environmental and ecological variables, which are themselves combinatorially vast and dynamic. Through this lens, the problem of building a predictive model of evolution appears impossible, stifled by biology’s curse of dimensionality.

Despite this challenge, recent studies have hinted that there may be low-dimensional structure in genotype-phenotype maps (2, 15, 18, 30, 38, 40, 41, 51). In most of these studies, the fitness of a particular genotype is portrayed as linear function of *K* underlying, latent phenotypes

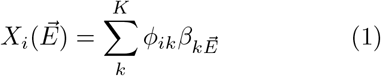

where, *ϕ*_*ik*_ is genotype *i*’s effect on latent phenotype *k*, and 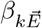 is the weight of that latent phenotype in environment 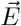.

These low-dimensional genotype-phenotype-fitness maps can be inferred via matrix factorization from large datasets composed of the fitness of many genotypes in many environments. This approach circumvents the difficulty of defining which phenotypes *should* be measured, and instead infers the underlying phenotypic structure from fitness variation. One study from our lab looked at a collection of experimentally-evolved adaptive yeast mutants and found that subtle environmental perturbations near the original evolution condition (Fig. 1a) were enough to induce variation in fitness that contained signals of the underlying latent phenotype space (18). We referred to these latent phenotypes, 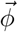, as “fitnotypes,” because they are fundamentally different from traditional, measurable phenotypes. Indeed, fitnotypes are only detectable as latent variables that contribute to fitness variation, while phenotypic changes that do not affect fitness are hidden. These fitnotypes, in turn, determine fitness based on environment-specific weightings. If adaptive mutants are functionally identical because they evolved towards the same fitness optimum, they should exhibit congruent behavior in fitness space regardless of the number of environments tested, and hence a one-dimensional version of Eq. (1) would suffice to capture fitness variation across multiple environments. Other studies have used a similar model to Eq. (1) to capture the fitness of thousands of QTLs from a yeast cross across 18 environments, or the behavior of human cell line knockouts under genotoxic stressors (31, 32, 38, 40), often including different constraints such as sparsity in the factorization problem.

**FIG. 1:**
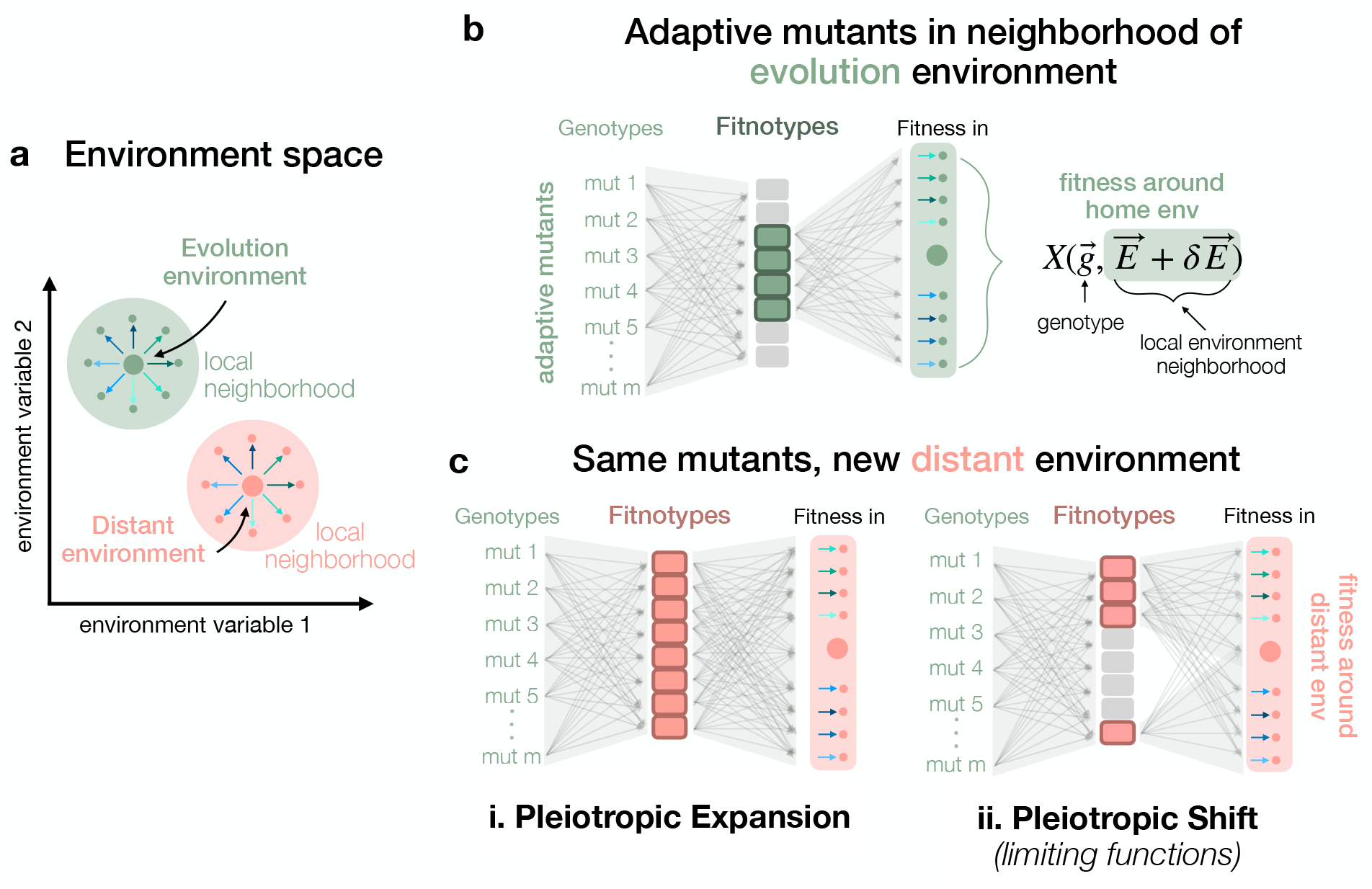
Two models for the nature of pleiotropy in adaptation. (a) Schematic of environmental structure in this study. Environments can be mapped onto a multidimensional environment space characterized by chemical and physical compositions. Large green dot represents an environment where adaptive mutants evolved (home base), and the large pink dot is a different “base” environment. Around each base, a set of identical environmental perturbations (arrows) is applied, generating clusters of similar environments around distinct base environments. (b) Schematic of fitnotype space for adaptive mutants near their home base environment. By measuring fitness in each of the green environments, we can infer how many fitnotypes matter for this set of mutants in their home environment. Here, only four of the possible 8 fitnotypes matter. (c) When we move the mutants to the distant base environment, and measure their fitness in all pink environments (base and perturbations), there are two possibilities. Either more fitnotypes become important and the space appears higher-dimensional (left, pleiotropic expansion), or the set of fitnotypes that matters shifts (right, pleiotropic shift).

All together, these studies suggest that low-dimensional structure underlies genotype-phenotype-fitness mapping across environments. But the source of this low-dimensional structure remains unclear, particularly in the case of adaptive mutants in subtle environmental perturbations (18).

One explanation is the “pleiotropic expansion” model. In this framework, mutations selected in one environment (Fig. 1a, green environments) are initially constrained to a low-dimensional phenotypic space, where only a subset of phenotypic effects matters to fitness (Fig. 1b, top). However, when placed in a new base environment with the same perturbations as above (Fig. 1a, pink environments), their pleiotropic effects become unconstrained, leading to an expansion in the number of phenotypes that influence fitness (Fig. 1b, bottom left). If true, this suggests that evolution dynamically constrains mutants to explore the “flattest” paths in phenotype space within a given environment, eliminating the costs associated with affecting too many phenotypes (34). Yet, in novel environments, no such constraints exist, allowing the full phenotypic diversity (and likely the associated costs) of a mutation to be exposed. This model reconciles observed polygenic adaptation and widespread pleiotropy with the theoretical idea of the cost of complexity (7, 33, 34). Here, the source of low-dimensionality is short-term evolutionary dynamics, where ascertainment bias in phenotypes skews adaptive mutants to appear low-dimensional only in their home environment (19).

An alternative explanation is the “pleiotropic shift” model (Fig. 1b, bottom right), in which adaptive mutants always affect many phenotypes, but only a small subset of these phenotypes are relevant in any given environment. As environments change, the relative importance of different phenotypes shifts, reweighting their contributions to fitness. Under this model, genotype-phenotype-fitness maps are consistently low-dimensional, both in the home environment and in distant environments, because fitness is determined by a small number of dominant, environment-specific challenges. This model bears resemblance to previous work on linear pathway models and Liebig’s law of the minimum, which states that growth is limited by the first essential nutrient or metabolite to run out (6, 22, 27, 46, 47). In analogy to this work on limiting nutrients or limiting metabolites, we conceptualize this pleiotropic shift model as indicative of “limiting functions.” In this qualitative model, the set of functions that a cell must perform in order to thrive is large, but at any given time there will be a small number of functions that limit fitness. Thus, only a small number of limiting functions can be profitably modified in adaptation. Meanwhile the pleiotropic side-effects on non-limiting functions are effectively hidden from natural selection. Just as nutrients can shift from being in excess to being limited as the environment changes, so too can functions can shift between being non-limiting and limiting in new environmental contexts. Such a model has implications for evolution across multiple environmental epochs, and bears on the question: how likely is it that the important functions in one environment will be important in all environments? In this model, the source of low-dimensionality is long-term evolution, which has shaped physiology to respond to environments in a generically low-dimensional way.

To differentiate between these two scenarios, we must build distinct fitnotype spaces for at least two different “base” environments, as in Fig. 1a, wherein adaptive mutants are measured in environments around their home base (evolution environment), and some other distant base. By comparing two different latent fitnotype spaces, we can characterize how the environment modulates genotype-phenotype-fitness mapping (Fig. 1b). In this work, we used a set of adaptive *Saccharomyces cerevisiae* mutants from previous evolution experiments, with known origins. We selected the home base and two additional “dis-tant” base environments, chosen to modify growth cycle and salinity of the environment. To each base, we applied an identical set of environmental perturbations (changes in temperature, glucose concentration, etc). We used singular value decomposition (SVD) to infer three separate latent fitnotype spaces for each base environment from fitness variation due to the perturbations, and compared their dimensionality. Our findings ultimately reject the pleiotropic expansion model and strongly support the pleiotropic shift model. We interpret these results as evidence in favor of a “limiting functions” model of fitness. Environments impose a low-dimensional constraint on evolution, as each environment presents a limited number of dominant challenges that shape fitness outcomes. These results highlight the fundamental role of environmental context in shaping the genotype-phenotype-fitness landscape, providing new insights into how adaptive evolution navigates complex and fluctuating environments.

## I. RESULTS

### A. An experimental design to probe environmental modulation of fitnotype space

To test whether adaptive mutants undergo a pleiotropic expansion or shift when they are moved away from their home environment, we designed an experiment to infer fitnotypes (latent phenotypic dimensions) for adaptive mutants in different “base” environments. In this study, we compiled a library of adaptive, DNA barcoded *Saccharomyces cerevisiae* mutants from previous evolution experiments (1, 19, 25). These mutants evolved in the same glucose limited media, under a serial dilution growth regime portrayed in Fig. 1a, but they differ in the time between passages (*T*_*transfer*_). Previous work has shown that fitnotypes can be inferred from fitness variation due to subtle perturbations around a base environment, so we chose three “base” environments around which to build our fitnotype maps: the two evolution conditions (*T*_*transfer*_ = 48 hours: “2 Day,” and *T*_*transfer*_ = 24 hours: “1 Day”), along with one more extreme environment, “Salt,” which has 0.5 M NaCl added to the glucose limited media and a transfer time of 48 hours. We applied a set of roughly 20 environmental perturbations to each base environment (Fig. 2b columns represent different perturbations on three base environments). These changes included small alterations to incubation temperature (30^*°*^C +/-2^*°*^C), in glucose concentration (1.5%+/-0.2%), or introducing sub-inhibitory amounts of drugs. See supplemental information for more details about the environments.

**FIG. 2:**
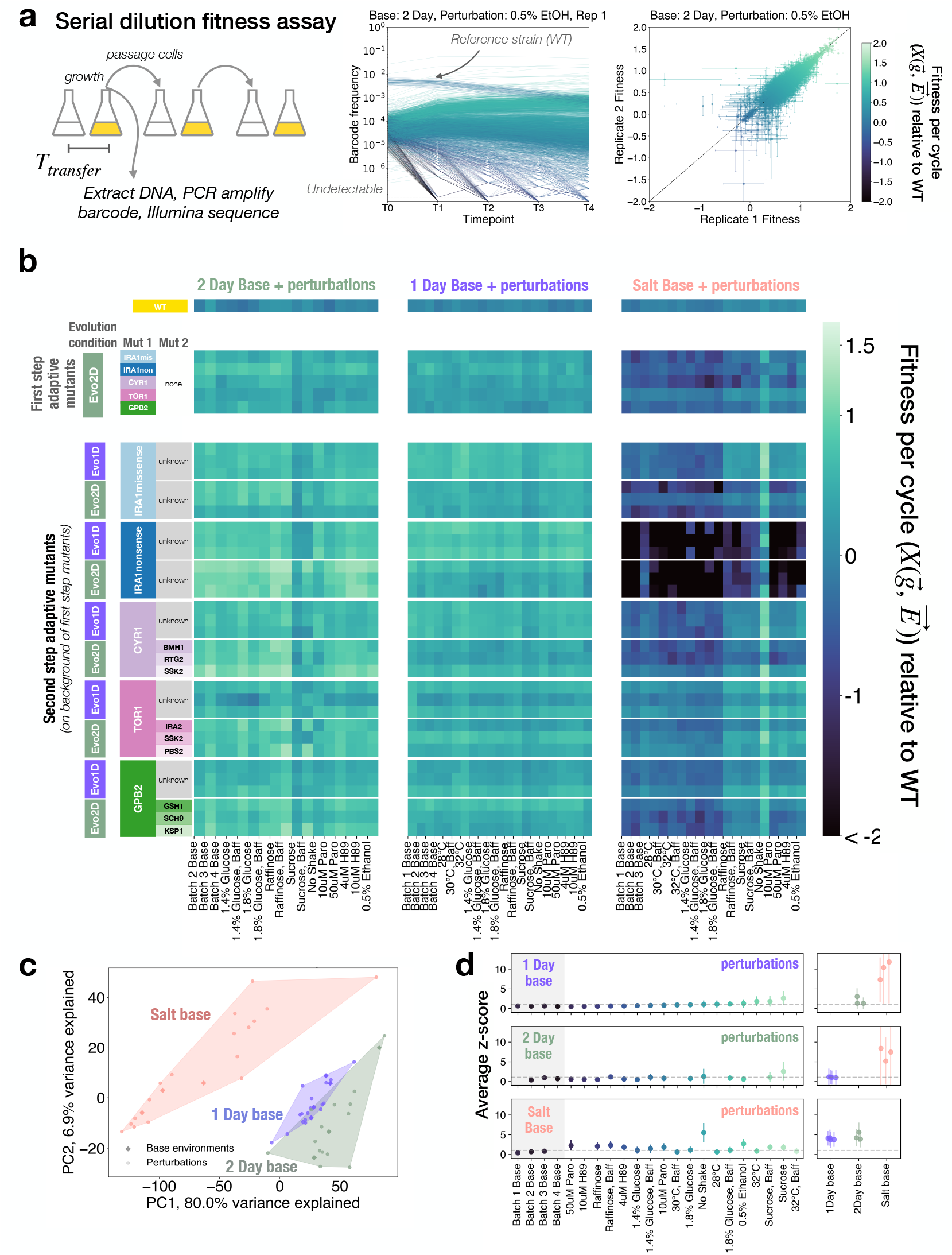
Fitness assay of adaptive yeast mutants in three base environments with subtle perturbations. (a) Fitness Assay Schematic: Barcoded yeast strains (4000) with adaptive mutations from previous evolution experiments are pooled with WT ancestor (95% of pool). The mixture undergoes serial dilution every 24-48h (T_*transfer*_). At each transfer, cells are preserved for DNA extraction, barcode amplification, and Illumina sequencing. This generates precise strain-specific frequency trajectories over time, enabling high-resolution fitness measurements relative to the ancestor (as shown by biological replicates) (b) Heatmap displaying genotype-by-environment fitness matrix. Colors indicate fitness values, 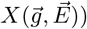. Columns represent environments (grouped by base), rows represent three mutants from each ancestor-evolution combination. Top row shows ancestral wild type, subsequent rows show strains with 1-2 additional mutations from either 1 Day “Evo1D”) or 2 Day (“Evo2D”) evolution environments.(c) PCA of fitness environmental covariance matrix. Environments projected onto top two PCs, with unperturbed base environments as solid diamonds. Convex hulls outline all environments within each base. (d) Average z-scores per environment, calculated from mutants’ fitness statistics across un-perturbed base replicates. Perturbation colors follow 1 Day base environment’s ascending z-score order. Dashed line indicates z-score=1. Right panels show other base environments’ z-scores relative to focal base.

To measure the relative fitness of all mutants in the library in a single environment, we utilize previously established methods that leverage DNA barcoding and high-resolution lineage tracking. We compete a pool of 4000 barcoded strains against a highly-abundant reference strain, the wild type (WT) ancestor, which takes up 95% of the culture. We quantify the frequencies of barcodes over time via amplicon sequencing. Based on these frequency trajectories, we can infer the relative fitness of thousands of strains. Fig. 1a shows the workflow from serial dilution experiments to highly replicable fitness inference (5, 18, 48). See Methods for more details on the fitness assay.

Fig. 2b shows a representative subset of our data, where each row is a mutant and each column is an environment. The mutants are organized by their background genotype and evolution environment, or home base, and the environments are organized by base. The most obvious signal comes from the tradeoffs present in Salt conditions, where mutants that are adaptive in 1 Day or 2 Day are much more likely to be deleterious with respect to the ancestor in the Salt base. This genotype-by-environment fitness matrix will be the basis of our further analyses.

First, we characterized each environment as an *m*-dimensional fitness vector, where *m* is the total number of mutants. This mutant-embedding of environment space is helpful because it allows us to focus solely on environmental differences that matter to fitness for our focal set of adaptive mutations. We used principal component analysis (PCA) to project each environment vector into a two-dimensional space, onto the first two PCs. In Fig. 2c, each point is one environment (unique base-perturbation combination). We colored each environment according to its base, and found that each set of perturbations belonging to the same base remains mostly localized in this two-dimensional space. This is especially informative because the first two principal components explain nearly 82% of the variance in the data. 1 Day and 2 Day are more similar bases to each other, as was visually evident in the heatmap in Fig. 2b, and Salt is a more distinct base. Therefore, our set of environments provides a range of environmental scales across which we can compare fitnotype maps of adaptive mutants.

Due to the large number of environments in our assay, we performed fitness measurements in batches of different perturbations, each containing one “un-perturbed” base, lending us batch replicates (diamonds in Fig. 2c). These serve as a natural baseline of variation, which we can leverage to evaluate the magnitude of our environmental perturbations. Differences between batch replicates reflect environmental factors that cannot be fully controlled experimentally, such as minor fluctuations in temperature and humidity, small inconsistencies in how the growth media were prepared, and biological noise (20). For each mutant, we calculated an absolute z-score in each perturbed environment compared to the batch replicates (see Methods). We took the average z-score across all mutants for that environment as a measure of the magnitude of the environmental perturbations, as “perceived” by the yeast cells and reflected in their fitness effects. Fig. 2d demonstrates that most perturbations within a base environment typically result in average absolute z-scores around 1 when compared to their batch replicates, indicating their very subtle effects on fitness.

All together, this experiment lets us compare fitness variation of distinct groups of mutants due to identical environmental perturbations, applied to different base environments. Conveniently, we have groups of mutants that each evolved to different bases, lending us more power to dissect the role of short-term evolution in shaping the fitnotype space we detect, and to differentiate between pleiotropic expansion and pleiotropic shift when the mutants are moved to a distant base environment (Fig. 1b).

### B. Environmental perturbations generate base environment-dependent fitness effects, revealing ExE interactions

While Fig. 2d suggests that the magnitudes of the same perturbation on different bases were similar, the “directions” might be different. For example, does adding 0.5% extra glucose to the media have a similar effect in a 24 hour dilution cycle and a 48 hour dilution cycle? And does the presence of salt disrupt the effect of additional glucose, if osmotic pressure exerts a more urgent challenge for the cells to respond to? The presence of environment-by-environment interactions might indicate that fitness in different base environments is dominated by distinct challenges, lending support for the limiting functions model of fitness.

To isolate a perturbation’s fitness effect on a particular mutant, we compute

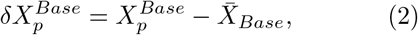

the mutant’s relative fitness in the perturbed environment minus its mean fitness across all batch replicates of the corresponding base. Fig. 3b shows *δX* due to the same perturbation on different base environments, for one mutant with a mutation in the IRA1 gene, evolved in the 2 Day home base from the WT ancestor. This mutant’s fitness relative to the WT is shown in the top right inset panel for batch replicates of each base environment, representing the baseline from which 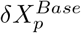 is calculated. Most perturbations show smaller fitness effects than the baseline mutation fitness effect, consistent with Fig. 2c, with several notable exceptions. The “No Shake” perturbation has little effect in the 1 Day and 2 Day bases, but has a highly beneficial effect on the relative fitness of this mutant in the Salt base, suggesting an interaction between salinity and oxygenation. The addition of 0.5% ethanol is also beneficial on a Salt background, but deleterious on a 1 Day background, and nearly neutral on a 2 Day background.

**FIG. 3:**
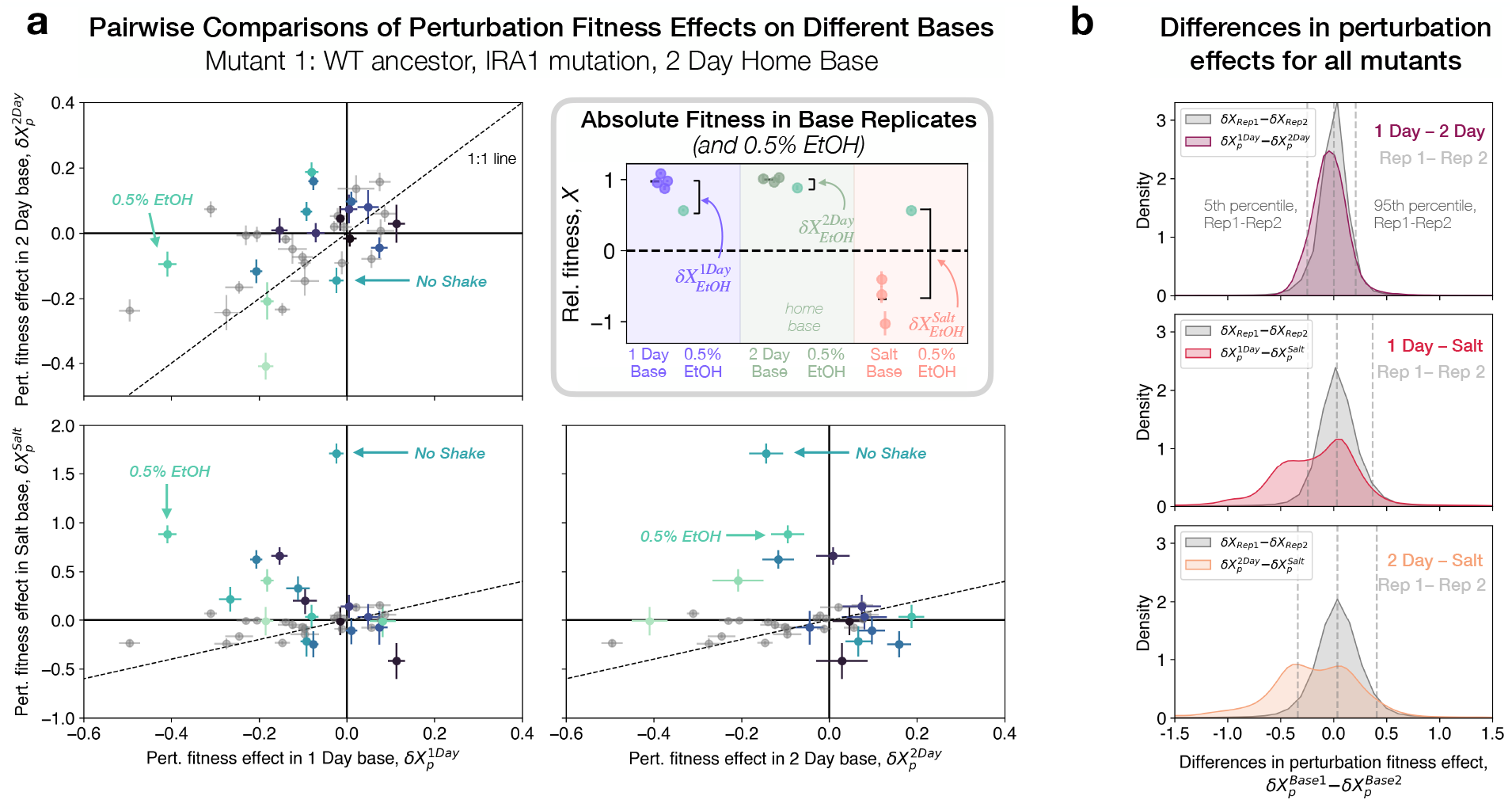
Widespread environment-by-environment interactions hint at phenotypic novelty. (a)Effect of environmental perturbation on this IRA1 mutant’s fitness across different bases. Each perturbation is a point, and the gray points compare *δX* for two replicates of the same environment. Perturbations that deviate from the 1:1 line more than noise would predict indicate the presence of ExE interactions. Inset: This IRA1 mutant’s (evolved in 2 Day base) raw fitness in each base environment, and in 0.5% EtOH perturbation. 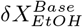 is difference between fitness in perturbation and base. (b) Difference in perturbation effects across different pairwise combinations of bases, aggregated across all mutants. In gray, the distribution represents noise-driven differences, obtained from replicate-replicate *δX* deviations. Vertical dashed lines represent 5th and 95th percentiles of the gray distribution. Colored distributions show pairwise comparisons of bases. 1 Day - 2 Day comparison (top panel) shows a small, but significant difference between the two distributions. Both comparisons with Salt (middle and bottom panels) have substantial density outside the gray distribution, suggesting the presence of many ExE interactions between Salt and the perturbations for many mutants.

Technical noise in the fitness measurement process can also be a source of uncorrelated *δX*_*p*_ across different bases, so we compared these base-to-base correlations with replicate-replicate *δX*_*p*_ correlations. We took two biological replicates from the same condition, and reasoned that any deviation from the 1:1 line could only be driven by noise, and could serve as a baseline for establishing a signal of ExE interactions. These replicate-replicate correlations are plotted in Fig. 3a as gray points (Pearson’s r=0.58, p=0.009). We find that for this mutant in IRA1, many perturbations exceed the spread in replicate-replicate comparisons, particularly in the Salt base, confirming that apparent non-additivity of perturbations is driven by more than just measurement noise.

We next aggregated the deviation from additivity (residuals from 1:1 line) for all mutants in all environments and compared them to this null expectation. In Fig. 3b, we plot the distribution of deviations from the 1:1 line for all pairs of biological replicates from the relevant bases in gray, which is narrow and centered around 0, compared to deviations across different base comparisons. The 1 Day-2 Day comparison shows a small, but significant (*KS* = 0.17, *p <* 10^−16^) difference in distribution from the replicate-replicate comparison, with 17% of comparisons demonstrating ExE interactions (FDR=10%). The 1 Day-Salt (*KS* = 0.37, *p <* 10^−16^) and 2 Day-Salt (*KS* = 0.37, *p <* 10^−16^) comparisons are bi-modal, with one peak corresponding to the technical noise-induced deviations, and another corresponding to significant ExE interactions across the two background bases. Here, 47% and 44% of comparisons, respectively, deviate significantly from the 1:1 line (FDR=10%). Hence, we see strong evidence for individual mutants behaving unpredictably upon the “same” environmental perturbations. In the supplemental information, we explore the correlation in *δX* for all mutants between pairs of environments, and find that certain perturbations show strong signs of ExE interactions. Overall, this suggests that genotype-by-environment-by-environment interactions may be crucial in shaping the fitnotypes we observe in each base environment.

### C. Emergence of GxExE interactions in low-dimensional fitness landscapes

These environment-by-environment interactions naturally emerge in a simple model of a fitness landscape, motivated by empirical observations of low-dimensionality across environments (18, 38, 40). To see this, we assume that fitness is mediated through a low-dimensional space of coarse-grained environmental variables, *ψ*_*k*_, where *k* = 1, 2, …, *K*. Therefore, we write the fitness of mutant *i* as a function of both its genotype, 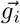, and the coarse-grained environmental variables 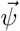.

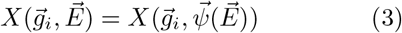

We model the fitness effect of an environmental perturbation by decomposing the environment into two parts: the base environment, *E*_*B*_, and the change in the environment due to the perturbation, *δE*_*p*_. If the perturbation is small, we expand the fitness function to write the change in fitness due to an environmental perturbation as

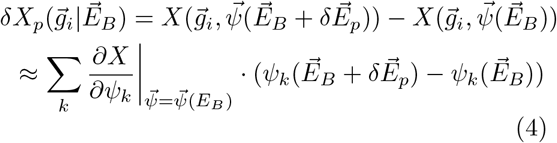

This equation, like Eq. (1), is a generalized version of the standard linear fitness models proposed in quantitative genetics literature, as well as an extension of the model used in Kinsler et al. to assign fitness to adaptive mutants via a low-dimensional latent phenotype space (18, 23). We can cast this equation for *δX*_*P*_ into more standard notation, where the partial derivative plays the role of a “trait” or a “phenotype,” and the 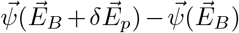 term is a selection gradient.

To simplify our notation, we can write

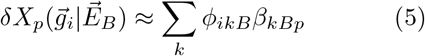

where *ϕ*_*ikB*_ is an effective phenotype that depends on the genotype and base environment, and *β*_*kBp*_ is the selection gradient that depends on the base environment and perturbation. Equation (4) shows that we should generically expect gxExE effects via the *δX*_*p*_ metric, unless the base environments are so close to each other that the 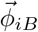 is actually base-independent, and the weights 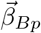 are the same across bases. Thus, the observations in Fig. 3 of ExE interactions are consistent with this simple model, since we chose our base environments to be far from one another. We can infer that they are distant enough to induce widespread gxExE interactions. Even so, if the environmental *perturbations* are small, the linear approximation in Eq. (5) is still a good one. We thus cull the Salt + No Shake perturbation (which was not subtle according to Fig. 2d), and henceforth use a linear fitness model as in Eg. (5) to probe the quantitative properties of environmental dependence of phenotypic landscapes. In particular, we can use Eq. (5) to infer *K* for each base environment, allowing us to differentiate between pleiotropic expansion and pleiotropic shift. Additionally, we can use this model to determine whether ExE interactions arise from a unique set of 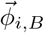 vectors in each base, or whether the weights 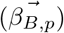 shift, shedding light on how limiting functions that determine fitness might change across different base environments.

### D. Environmental shifts away from home base do not induce pleiotropic expansion

Having established that environment-by-environment interactions are prevalent in our data, we next sought to distinguish whether adaptive mutants fit the pleiotropic expansion or pleiotropic shift models presented in Fig. 1, using a linear model of fitness as in Eq. (5).Kinsler et al. proposed that adaptive mutations could have large fitness effects *and* be pleiotropic because they are modular near their evolution condition(18). In other words, a set of adaptive mutations might look the same in environments that are very similar to their evolution condition because they affect fitnotypes in the same way. But there are no guarantees for low-dimensional behavior amongst this set of adaptive mutants when their fitness is measured in new environments that are dissimilar from the evolution condition. Under this model, we would expect the fitness of a set of adaptive mutants to be well-approximated by a low-dimensional model in the home base (and subtle perturbations). But as the environment shifts, new traits may become important for survival, so the previously hidden phenotypic effects may reveal themselves. Does this manifestation of hidden pleiotropy tend to drive higher-dimensional behavior of adaptive mutants in foreign environments?

To test this, we first focused on the adaptive mutants that arose on the WT background in the 2 Day base environment. We partitioned our environments by base and asked whether the same set of mutants yielded the same low-dimensional behavior around each base environment. Specifically, we wondered whether the 2 Day base would look different from the other two bases because it has the special distinction of being the evolution environment for this set of mutants. Under the “pleiotropic expansion” model, we expect the dimensionality of the mutant set to appear higher around Salt and 1 Day than 2 Day.

To infer the dimensionality of the latent space, we did singular value decomposition (SVD) on the *δX* matrix of perturbation effects on fitness in each base separately, and quantified the variance explained by each dimension. By definition, the first *k* SVD components generate the *k*-dimensional linear model with the lowest reconstruction error, and as *k* approaches the rank of the input matrix, the variance explained approaches 100% (11).

We compared the fraction of variance explained by each component across the three different base environments. Due to different levels of noise and measurement resolution, we have different limits of detection for each base, though the 1 Day and 2 Day limits of detection nearly coincide. We compared the number of components that fall above the variance explained by the limit of detection. We obtained this limit by running SVD on 1000 folds of a noise-only matrix, filled with 0-centered Gaussian random variables, with standard deviation equal to inferred standard error due to measurement noise in the fitness assay. As shown in Fig. 4a, the *δX* of the mutants that evolved in the 2 Day base are well-approximated with a 4-dimensional model in environments belonging to the 2 Day base. When the same set of mutants is measured in the same set of environmental perturbations on the 1 Day base, a 3 components fall above the limit of detection. And the most distant environment, Salt, also reveals 3 dimensions. So for this particular set of mutants, we find that dimensionality in the evolution condition base, 2 Day, does not substantially differ from the non-evolution condition bases, and in fact is slightly higher.

**FIG. 4:**
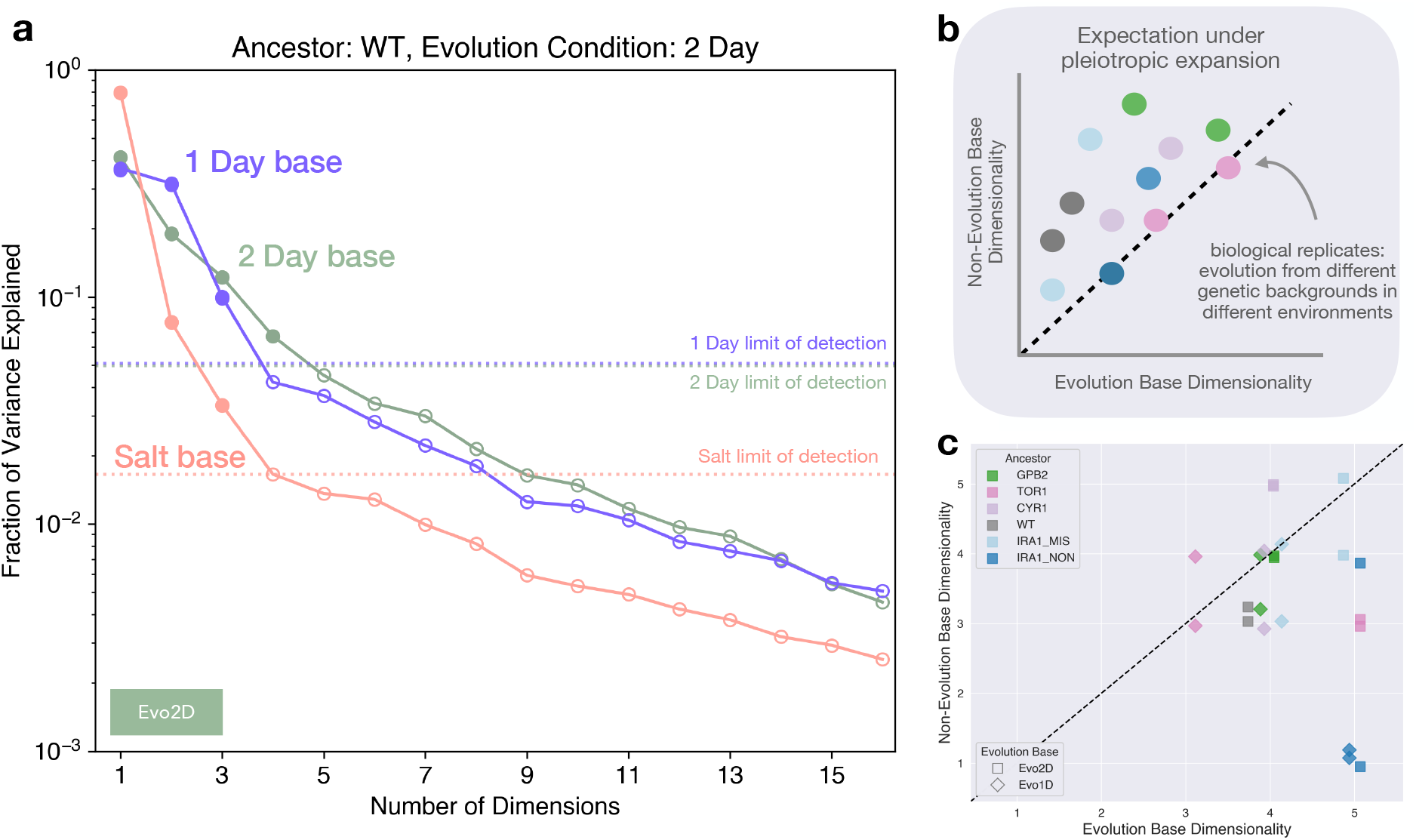
Pleiotropic expansion model is not supported by dimensionality reduction across all bases. (a) Fraction of variance in *δX* explained by component for each of the three environmental bases. Points are colored if they fall above limit of detection. Limit of detection was obtained by doing SVD on 1000-folds of noise-only matrices, using inferred measurement error as the standard deviation for the random variable in each fold. These mutants evolved in 2 Day base, and have 4 dimensions above the limit of detection in 2Day, 4 in Salt, and 3 in 1Day. (b) If pleiotropic expansion were true, we should expect systematically higher dimensionality in a non-evolution condition base than an evolution condition, amongst a set of biological replicates (ie independent evolution experiments). (c) Dimensionality in evolution environment and non-evolution environments for different “biological replicates” - sets of evolved mutants from different ancestral backgrounds, and in different environments (jitter added). We find no evidence for pleiotropic expansion.

But this set of mutants effectively yields only two comparisons – home base with two other bases. To fill out this picture, we leverage the second-step mutants from subsequent evolution experiments, represented as lower rows in Fig. 2b. These mutations arose in both the 2 Day base and 1 Day base, so we now have two evolution conditions to compare, and to lend more evidence for or against the pleiotropic expansion hypothesis. These sets of mutants serve as biological replicates – they represent multiple instances of diversification from different genetic backgrounds, in two different environments. For each set of mutants, we can quantify their dimensionality in their evolution environment, and the other two bases. We visualize these comparisons as a scatter plot, with dimensionality in evolution condition on the x-axis, and dimensionality in distant environmental base on the y-axis (Fig. 4b). This panel shows our expectation under the pleiotropic expansion model, whereby dimensionality in the non-evolution base is systematically higher than the evolution base, for the same set of mutants. Fig. 4c shows the inferred dimensionality for all genetic backgrounds and evolution conditions in this study. Each point is a set of adaptive mutants, whose dimensionality is quantified using the approach from 4a. We include the scree plots of fraction variance explained by number dimensions for all other mutants in the supplemental information.

Overall, we find no evidence for the pleiotropic expansion hypothesis. In general, there is no clear relationship between dimensionality and whether a group of mutants evolved to the base or not. In fact, the scatter plots indicate that the evolution condition appears to be higher-dimensional more often than the other bases, but more replicates would be necessary to confidently make such a statement. We can claim, however, that a similar number of dimensions is necessary to build fitnotype maps for each set of mutants, regardless of where they evolved. We explore several other methods to infer dimensionality in the supplemental information. Additionally, we quantify the dependence of inferred dimensionality on number of environments included in the input matrix in the supplemental information. None of these additional analyses change our conclusions. This allows us to reject the pleiotropic expansion hypothesis, and turn our attention to quantifying how fitnotype space shifts across distinct base environments. In the rest of this work, we aim to characterize how different base-dependent limiting functions influence the way fitnotypes in different base environments map onto each other.

### E. Characterizing pleiotropic shift across base environments via fitness prediction

We have shown that each base environment reveals a similar number of fitnotypes for all sets of mutants, suggesting that low-dimensionality is not specific to the evolution condition. We next wondered the extent to which different fitnotype spaces overlap. Are these base environments limited by the same functions?

It is difficult to ask this question directly because fitnotypes are latent variables, so instead we use predictive power as a proxy for fitnotype overlap. Prediction loss is invariant to the choice of basis in matrix factorization, so it is a useful metric for this task. Fig 5a presents our approach. First, we select one base as the “training base,” and hold out a single perturbation *i*. We do SVD on the training base (minus *i*) to discover fitnotypes for this base (green boxes) for all mutants. Then, we use these as features in a linear regression model, and fit new “weights” on the fitnotypes to predict *δX* in the held-out perturbation *i*, obtaining a leave-one-out cross-validation error. Similarly, we can fit weights on the green fitnotypes for any perturbation, including those from other bases, yielding prediction error for “test base” perturbations. We aggregate prediction error for all held-out perturbations within the training base, and all perturbations from the other two test bases, to quantify the cross-validation predictive power of each fitnotype to compare. Importantly, the cross-validation error even within the “training” base is still a test error, because that perturbation was left out of the initial fitnotype discovery. We are essentially asking: if we allow our model to freely reweigh training fitnotypes to best predict *δX* in a new base, how close can we get to within-base predictability?

**FIG. 5:**
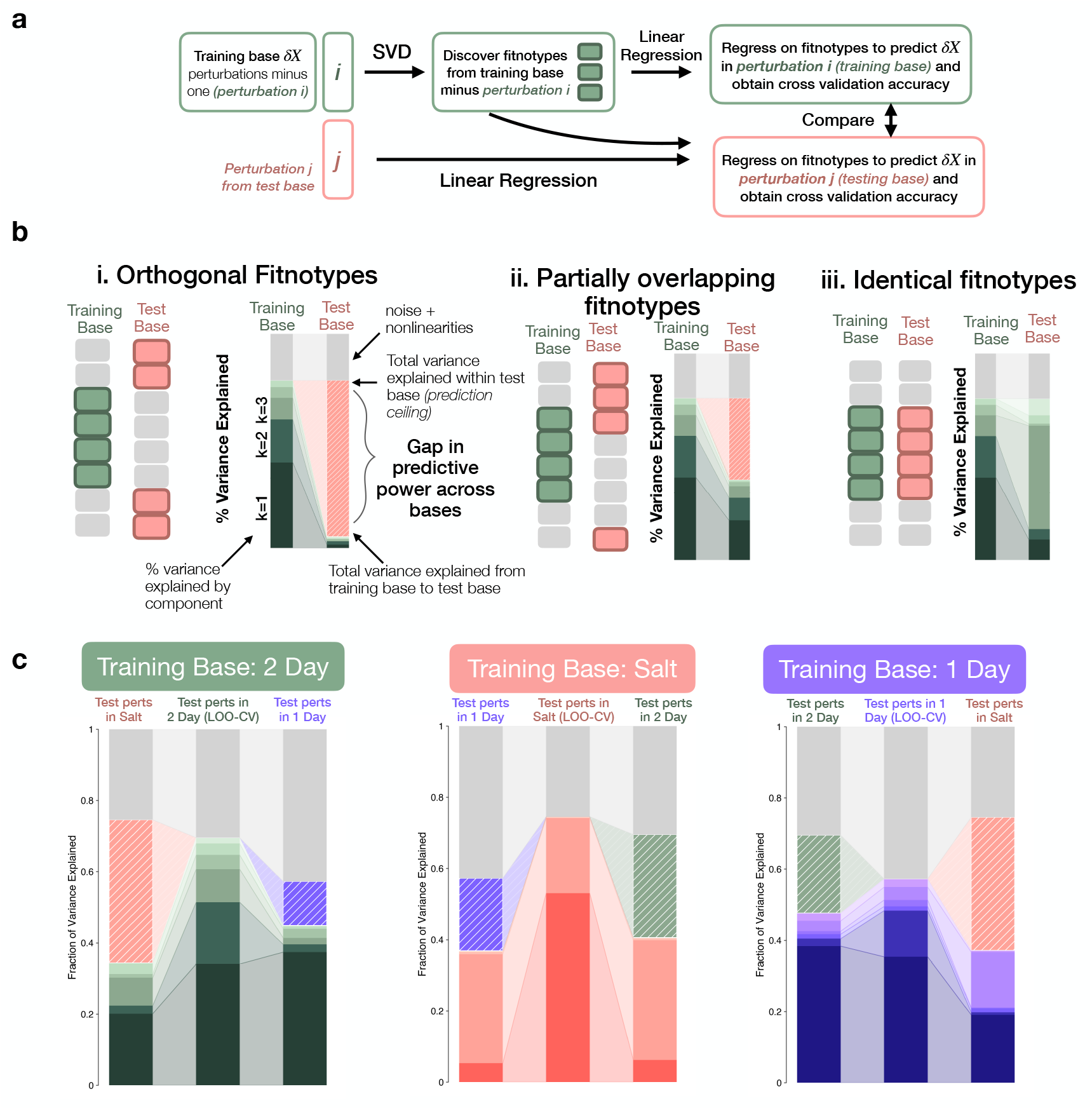
Predicting perturbation fitness effects within and across environments reveals structure of limiting functions. (a) Prediction workflow: We partition training data to extract base-specific fitnotypes via SVD, use these in linear regression to predict fitness changes in held-out perturbations (*δX*), and quantify variance explained within and across environments. (b) Expected prediction patterns in different scenarios: (i) orthogonal limiting functions between bases prevent cross-base prediction; (ii) partially overlapping functions allow some cross-base prediction with characteristic gaps (pink dashed section is missing test base variance, gray is unpredictable variance); (iii) identical functions yield equivalent predictive power across bases. (c) Our choice of mutants and base environments reveal partially overlapping fitnotypes, supporting scenario (ii).

We have different expectations for this predictive power across bases under different models of how fitnotype spaces map onto one another, as shown in Fig. 5b. First, we note that even within the training base there will be unexplained variance due to noise and/or nonlinearities that cannot be captured by a 6-dimensional linear model. We represent this unexplained variance in gray. If fitnotypes are fully orthogonal (Fig. 5b, i.), we do not expect that any amount of re-weighting will allow us to fit *δX* in the testing base, so the total variance explained by all training fitnotypes (green bars) will be negligible in the test base. We indicate the maximum achievable prediction had we inferred fitnotypes directly from the test base with a “prediction ceiling,” and highlight the gap in predictive power using the test base color and diagonal white lines. This scenario corresponds to a strong limiting functions model, where fitness-relevant functions are entirely distinct across environments and *ϕ* vectors are orthogonal, clearly yielding ExE interactions (Results C, Eq. (5)). At the other extreme (Fig. 5b, iii), fitnotypes fully overlap between bases. Although weights (*β*) may differ, the underlying functions remain the same, leading to comparable predictive power across bases because linear regression allows the flexibility of re-weighting. This still reflects ExE interactions, but they are driven by changing weights rather than fitnotypes. Between these extremes (Fig. 5b, ii), environments may harbor some of the same fitnotypes, but not all. These shared limiting functions yield moderate predictive power, but the distinct functions also create a substantial gap in predictability between the training and test bases. We confirm these expectations with simulated fitness data in the supplemental information.

In Fig. 5c, we compute the variance explained by different fitnotypes as presented above, for all mutants. We find partially overlapping fitnotypes between all base environments. As anticipated, the limiting functions between 1 Day and 2 Day appear to be more similar, as the gap in prediction power between 2 day and 1 Day is smaller than between 2 Day and Salt (Fig. 5c, left panel). We also observe some reweighting of fitnotypes in different bases. For example, the fifth 1 Day fitnotype contributes little to predict *δX* in the training base, but grows to explain nearly 20% of the variance in Salt (Fig. 5c, right panel). Note too that we fit a 6 dimensional model in the linear regression (Fig. 5a) to lend extra flexibility, but we see evidence for much lower dimensionality, particularly in Salt, which is dominated by just two fitnotypes. In the supplemental information, we show similar prediction analyses for individual sets of mutants, and predictive power for individual test perturbations within the bases. We also show in the supplemental information that a genotype-fitness model that includes fitnotypes and flexible re-weighting outperforms other low-dimensional models (i.e. simple gene-level predictions) for predicting *δX*.

Taken together, these findings combine to strongly support the pleiotropic shift hypothesis, with partially overlapping fitnotypes across different bases. We see direct evidence for phenotypic variation that remains hidden in certain bases, but that is revealed in others. Additionally, within the overlapping phenotypic space, the traits that most strongly contribute to local fitness often shift. It is worth noting that the inputs to our model for each base environment are, on the surface, identical. The same genotypes and the same physical and chemical changes to the base environment make up the rows and columns of the input matrix. Therefore, the heterogeneity in fitnotype space across base environments is entirely attributable to how the base environment constrains the fitness function. This lends support for the idea that fitness in a given environment is dominated by a small number of limiting functions, and highlights the importance of environmental context in the inference of genotype-phentoype-fitness maps.

## II. DISCUSSION

In this study, we aimed to understand how genotype-phenotype-fitness maps shift across environments. Building such maps has been a difficult task for several reasons. (i) Genotype space is vast, (ii) it is unclear which phenotypes are important to fitness, and (iii) environments can modulate both the genotype-to-phenotype mapping, and phenotype-to-fitness mapping. Previous work tackled the initial question of mapping phenotype space for a group of adaptive mutants near their evolution environment, using a top-down linear model of fitness variation across subtly perturbed environments (18). This technique yields a map of abstract “phenotypes” that are revealed through their effect on fitness that Kinsler et al termed “fitnotypes.” In generating full genotype-fitnotype-fitness maps around distinct base environments, which are themselves distant from each other, we were able to explore the nature of local and global pleiotropy for adaptive mutants and distinguish between two models of how pleiotropy might manifest itself.

One hypothesis, the pleiotropic expansion hypothesis (Fig. 1a), is that the dimensionality of a model capturing the fitness variation of adaptive mutants is small near their evolution condition, but grows dramatically around distant bases. We did not see evidence supporting this model: we reliably found instead that the dimensionality of our models were consistently low across bases, regardless of whether they evolved in the base environment or not. This allowed us to reject the “pleiotropic expansion” model.

We instead found strong evidence for the alternative, the pleiotropic shift model (Fig. 1b). In this model, genetic mutations do generate a pleiotropic collection of phenotypic effects, but the dimensionality of the phenotypic variation that affects fitness remains limited even in environments that are distant from the evolution condition. This pattern is consistent with a limiting functions model of fitness. In any particular environment, there is a small number of key limiting functions that can be altered to affect fitness. These are the functions that determine the cells’ ability to surmount limiting challenges and thrive. Fitness variation is thus dominated by this small set of functions, suggesting that phenotypic diversity in an adapting population is constantly being projected down onto a few key dimensions. The choice of these dimensions changes as the environment shifts.

How might such a pleiotropic architecture of limiting functions arise? Similar questions have been explored extensively in the context of nutrient availability (22, 27, 47). If cellular growth is a function of available nutrients, many different nutrients may be required. However, at any given time, growth will be constrained by just one of these—the limiting nutrient—while the others remain in excess. This principle is well understood in both ecology and chemistry; for example, in a chemical reaction, the limiting reagent determines the reaction’s progress. Fitness is a more complex function than a simple chemical reaction, but at its core, it is governed by a series of biochemical processes that drive growth. Differences in these processes create differences among cellular performances, manifesting as fitness advantages or defects. It is therefore reasonable to extend the concept of limiting nutrients or reactants to the level of fitness. Importantly, the key constraint on fitness may not always be nutrient deficiency. It could be the need to withstand extreme heat, buffer against extreme pH, or overcome some other environmental challenge. Ultimately, a cell “per-ceives” its environment as a set of challenges, but in any given moment, closely related cells will be limited by only one or a few of these challenges. After all, optimizing growth rate is irrelevant if the cell is dying from osmotic stress.

We are presented with a range of possibilities for how environments might vary under this model. Two distant environments might present entirely orthogonal selection pressures to a population. In a simple case, one environment might have an abundance of a fermentable carbon source for the yeast, so optimizing growth rate or lowering lag phase for fermentation metabolism the best function to optimize in a fitness competition. But a different environment might only have non-fermentable carbon sources, and so changes to fermentation metabolism will have little effect on fitness in this second environment. Any pleiotropic effects from the first environment that bear on respiration metabolism will be revealed, and differentiate between the various mutants. Imagine another scenario, in which both of these media are placed in an incubator at 37^*°*^C. Now, while the base media are still “orthogonal” metabolically, the yeast are most concerned with surviving the heat shock, and so the limiting function is heat survival in both environments. Any variation in that phenotype will dominate the relative fitness between mutants in both environments, so we will infer these underlying fitnotypes to be fully overlapping. Conceptualizing environments in this manner presents an intriguing possibility for understanding environmental similarity, which has historically been difficult to define. The notion of overlap in limiting functions could be a useful organism-specific metric for parameterizing environmental distance, from the perspective of competitive fitness.

One natural question to emerge is why there seems to be overlap in the phenotypes affected in our pool of mutants, and the limiting functions that are important in our distant, non-evolution base environments. A priori, it is not clear that variation in the hidden phenotypic effects from a group of mutants will necessarily be relevant to the limiting functions in an arbitrary environment.

The most obvious possibility is that perhaps the bases are not in fact sufficiently distant. All three are lab environments with glucose-limited media, so while we do observe many differences in fitness, perhaps we are constraining the environment space more than we thought. Or perhaps the very fact that yeast cells can thrive in all these environments itself necessarily acts as a built-in constraint on fitness assays and growth, meaning that the range of possible environments will always be limited. So it is interesting to speculate: what will we learn when we apply many more perturbations to a very large number of base environments? Perhaps we will discover that there are endless ways to project phenotypic diversity onto fitness, and that at a certain point the collection of genotypes will become the limiting variable because phenotypes are indeed low-dimensional (16, 21, 42). Or perhaps we will discover that environment space, constrained by survivability, is actually small, and consistently mappable. These extensions of environment space provide a promising avenue for future work.

Still another possibility for why we tend to detect limiting functions across the three base environments is that single bouts of evolution can exhaustively explore all phenotypic diversity, due to the tight integration of cellular and genetic networks, such that any possible important phenotype is already present in an adapting population of sufficient size (8, 14, 19, 49). Or, perhaps we are biased by our genotype set. Most of the mutations in our collection are either autodiploidization events, which affect the entire genome, or in signaling pathway, which are known to be especially pleiotropic (44, 48). These mutants are also atypical in that they harbor large beneficial fitness effects, despite their genetic proximity. What might we find by applying similar environmental perturbations and measuring the relative fitness of a different set of genotypes? Perhaps a library of “random” mutations from transposon insertion, for example, would generate more novel behaviors across environments. It would also be enlightening to compare our results to a set of genotypes that have adapted for much longer than ours, either from experimental evolution with the LTEE *E. coli* lines for example, or as a result of evolution in natural settings by looking at a large collection of natural isolates that represent the extent of standing genetic diversity within a species (13, 28, 39).

The limiting functions model naturally predicts that the fitness effects of environmental perturbations can often be heavily dependent on the base environment. If base environments present distinct challenges, the projection of fitness onto these key dimensions will be different, even as a result of the same underlying physical and chemical changes. Thus, idiosyncrasy in *δX* effects across bases is to be expected. However, this idiosyncrasy does not mean that prediction writ large is hopeless. The fact that dimensionality remains constrained locally in any base environment raises the possibility of practical applications, because the total space of fitness variation is smaller than we might expect *a priori*.

More broadly, the limiting functions model offers insight into how evolution might proceed over successive environmental epochs. First, a clonal population accumulates multiple adaptive mutations that compete with each other for fixation. All of these adaptive mutations confer a fitness benefit relative to the ancestor, and tend to affect similar phenotypes that matter to fitness in their environment, but they harbor a reservoir of hidden phenotypic diversity. When the environment shifts and a new epoch begins, a different set of phenotypes will be important for fitness, reshuffling the relative fitness of each mutant. The pool of phenotypic diversity amongst a group of adaptive mutations is large when integrated across multiple environments, but is never so large in a single epoch that mutants behave in completely idiosyncratic ways. Environments present limiting challenges that impose low-dimensional structure on phenotype space, and dominate the fitness response. This low-dimensionality provides some hope that evolutionary prediction is possible, and brings us one step closer to the ambitious goal of building a universally relevant genotype-phenotype-fitness map.

## III. METHODS

### A. Experimental procedure for barcoded fitness assay

#### 1. Experimental overview of fitness assay

We performed fitness assays as previously described (18, 26, 48). We pool all barcoded strains and compete them together against a reference strain (in this case the wild type ancestor, WT, represented at 95% of the pool at the beginning of the competition) by growing them for a predetermined transfer duration (*T*_transfer_) in 100 mL of glucose-limited media. We grew the cells in incubators set at temperatures determined by environmental prescription, for either 24 hours or 48 hours per cycle. All environments except “No Shake” were rotated at 223 rpm. At the end of each cycle, after the cells have grown in direct competition, we transfer 400 uL of saturated media into fresh media, or roughly 5 × 10^7^ cells, preferentially sampling lineages that have grown to higher frequency over the course of the cycle due to increased fitness relative to the mean population fitness. We dilute the cells at a 1:250 ratio, corresponding to roughly 8 generations of cell division over the course of the growth cycle, and we perform 4 total dilution cycles. At each transfer, we save the rest of the untransferred cells in sorbitol for DNA extraction and PCR amplification of the barcode region. We sequence 5 timepoints to get estimates for the frequency of each barcoded lineage, and use the log-fold change in frequency to infer the relative fitness of each strain with respect to the WT ancestor, correcting for the changing population mean fitness. With these methods, in each environment we were able to infer the relative fitness of thousands of strains. We perform fitness assays in multiple environments (Supplemental information).

#### 2. Yeast strains

All strains in this study are of genetic background Mat*α*, ura3Δ0::Gal-Cre-KanMX-1/2URA3-loxP-Barcode-1/2URA3-HygMX-lox66/71. Mutant strains come from the evolution experiments described in (1, 19, 25). The mutations that differentiate the barcoded strains from the WT arose in several evolution experiments via serial dilution. One study, Levy et al, generated an initial set of adaptive mutants from a single WT ancestor that evolved in glucose-limited media with 48 hour dilution cycles (*T*_transfer_ = 48 hr) over 160 generations (25). Each mutant typically harbors one putatively causal genetic variant that improves fitness relative to the ancestor in this environment. Most of these mutations were either auto-diploidization events, or single nucleotide changes in genes in the Ras/PKA and TOR/Sch9 pathways, both nutrient-sensing pathways (48). In subsequent studies, four of the Ras/PKA mutants and one TOR mutant were re-barcoded and further evolved in two environments: the original 48 condition (hereafter, “2 Day”) and a 24 hour condition (hereafter, “1 Day”)(1, 19). The cells complete fermentation of the glucose at around 20 hours and switch to non-fermentable carbon sources through respiration afterwards, so the 1 Day condition greatly limits the time the cells spend in respiration, amounting to a significant environmental shift despite unchanged media (26).

#### 3. Growth environments

All of our growth environments are based on M3 minimal, glucose-limited media. Our environments are defined by the media in the flask, additional components added to the media, the shape of the flask, the shaking status, and the temperature of the incubator. Due the scale of our experiment, we grouped our environmental perturbations into 4 batches, each with 3 base conditions and between 4 and 5 perturbations. This meant a total of 12-15 environments per batch. We had two biological replicates per environment. The first batch was temperature perturbations, the second batch was glucose gradients, the third batch was additional carbon sources, and the fourth batch was drugs and ethanol. See Supplemental Information for a full list of environments.

#### 4. DNA Extraction

We extracted genomic DNA according to a similar procedure in (18). Briefly, we thawed 400 *µ*L of cells for each sample. We spun down the cells at 3500 rpm for 3 minutes, and discarded the supernatant. We washed the cells by adding 400 *µ*L of sterile water and spun them down at 3500 rpm for 3 minutes, and discarded the supernatant. We re-suspended the pellet in 400 *µ*L of extraction buffer, and incubated at 37^*°*^C for 30 minutes. Then, we added 40 *µ*L of 0.5M EDTA, and vortexed. Then, we added 40 *µ*L of 10% SDS, and vortexed. We next added 56 *µ*L of proteinase K, and vortexed very briefly. We incubated at 37^*°*^C for 30 minutes, and put the samples on ice for 5 minutes. We then added 200 *µ*L of 5M potassium acetate, and manually shook the tubes. We incubated on ice for 30 minutes. Then, we spun the tubes for 10 minutes at maximum speed (13000 rpm), and transferred supernatant to a new tube with 750 *µ*L isopropnaol, and rested it on ice for 5 minutes. We spun it down for 10 minutes at max speed and discarded supernatant. We eashed twice with 750 *µ*L of 70% ethanol, vortexing very briefly and spinning at maximum speed ofr 2 minutes, then discarding liquid. We resuspended in 50 *µ*L Tris pH 7.5, and left it on the bench overnight if the pellet was not fully resuspended. We next added 1 *µ*L of 20 mg/mL RNase A and incubated at 65^*°*^C for 30 minutes. To digest the ancestor (so that we do not sequence our high-abundance reference strain), we added 5.5 *µ*L of Cutsmart buffer and 1*µ*L of ApaL1 restriction enzyme to each tube. We incubated the tubes at 37^*°*^C overnight, and quantified with Qubit.

#### 5. PCR amplification of barcode locus

To amplify the barcode region for sequencing, we perform two steps of PCR. For the first step, our goal is to tag individual molecules with UMIs and include first step primers to allow for combinatorial indexing. We divided each sample into 8 separate PCR tubes in order to eliminate the influence of PCR jackpotting. For a single set of 8 reactions, we used 200 *µ*L HotStartTaq Polymerase master mix, 8 *µ*L each of forward and reverse primers (10 *µ*M), 118 *µ*L nuclease-free water, 16 *µ*L of 50 mM MgCl2, and 50 *µ*L of genomic DNA. We ran 3 cycles of the following program.

1. 94^°^C 10 min
2. 94^°^C 3 min
3. 55^°^C 1 min
4. 68^°^C 1 min
5. Repeat steps 2-4 for a total of 3 cycles
6. 68^°^C 1 min
7. Hold at 4^°^C

We did a clean-up (standard DNA purification procedure) of the step 1 PCR product, and then performed step 2 PCR. We split our template DNA into 3 reactions, and for a set of 3 reactions, we used 45 *µ*L template, 65.5 *µ*L nuclease-free water, 30 *µ*L Q5 buffer, 2.5 *µ*L 10uM Forward and Reverse Nextra primers, 3 *µ*L 10mM dNTPs, and 1.5 *µ*L Q5 polymerase. We then ran the following program for the second step of PCR, which amplifies for 20 cycles.

1. 98^°^C 30 sec
2. 98^°^C 10 sec
3. 62^°^C 20 sec
4. 72^°^C 30 sec
5. Repeat steps 2-4 19 times
6. 72^°^C 3 min
7. Hold at 4^°^C

We did a sample clean-up using standard DNA purification techniques, and quantified with Qubit.

#### 6. Sample pooling and amplicon sequencing, and data processing

We pooled our samples and sent them to Admera Health for quality control, bead cleanup, and sequencing. We sequenced on HiSeq and Novaseq S4. We processed our data in exactly the same way as (18) to obtain barcode counts and frequencies for each sample, from raw sequencing reads.

### B. Analysis

#### 1. Fitness inference

We adapted the fitness inference method introduced in Ascensao et al (5), and closely follow their derivation of maximum likelihood fitness estimate below. We want to infer the fitness of a barcoded yeast strain from the read count information. Read counts for a particular barcode are generateed through a sequence of noisy processes, such as stochastic growth, bottle-necking, DNA extraction, PCR amplification, and sequencing. Each of these processes is individually a counting process, so we can model the read count of barcode *i* at timepoint *t* as a negative binomial random variable, which essentially allows us to think about the actual value of the read count as being sample from an over-dispersed Poisson distribution, where the variance is larger than the mean:

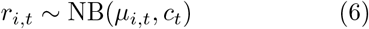

where the expected value of the read count is

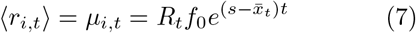

and the variance is

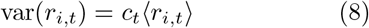

There are several parameters we need to infer before we can try to infer *s*. In order to calculate the expected value of the read count, ⟨*r*_*i,t*_⟩, we must have an estimate of the mean fitness of the barcoded population with respect to the ancestor, 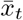, which we can infer from the trajectories of known neutral barcoded lineages. For a set of neutral barcodes, we have (by definition):

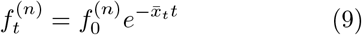

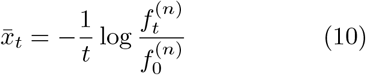

We can actually calculate this, with the caveat that read counts are somewhat noisy estimates of the underlying frequency. Also, since we assume all the barcodes that identify neutral lineages should have identical fitnesses (s=0), we can take the sum of all neutral barcodes at a given timepoint, 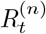, to calculate the mean fitness.

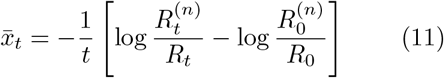

where

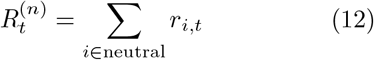

Alternatively, we can calculate the mean fitness within time intervals, where we have

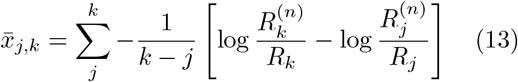

We can estimate the error in the mean fitness value by bootstrapping over neutral barcodes and recalculating the mean fitness. Alternatively, we can calculate the median absolute deviation between subsamples of neutral barcodes. To infer the variance of the negative binomial distribution, we can also leverage our neutral barcodes. We can think about all the potential sources of noise in the stages from cell number in the flask to read count. There is biological growth noise and bottlenecking noise, which should be correlated across timepoints (ie it accumulates), and there is uncorrelated technical noise from DNA extraction, 30 rounds of PCR, and sequencing. All of these processes are counting processes, so we expected the variance of the frequencies to be proportional to the mean. Note, we not are treating the total coverage at a particular timepoint as a random variable. So we have

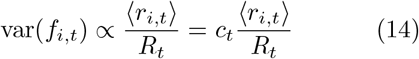

Given that we have a variance that is strongly dependent on the mean, we apply a variance-stabilizing transformation. Let’s define

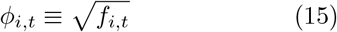

Applying the square root transformation effectively decouples the variance from the mean. We can think about how the variance in *f*_*i,t*_ is related to the variance in *ϕ*_*i,t*_. We can start with the Langevin equation for the change in frequency of a mutation due to drift.

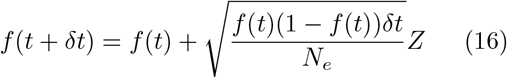

Consider the change in *ϕ* over one time interval:

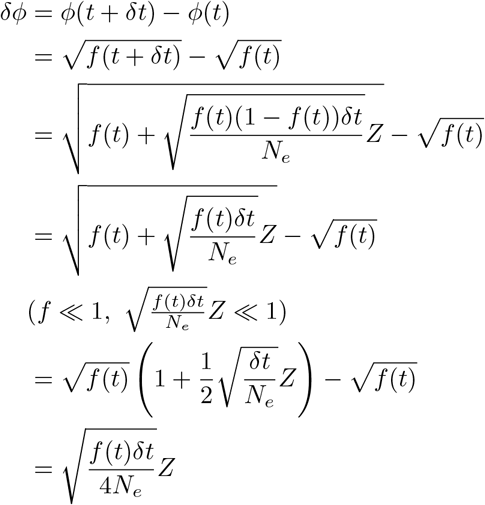

So the variance in *ϕ* due to genetic drift over a single cycle is 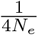. Between two timepoints, we expect neutral lineages to change due only to noise from genetic drift (which accumulates over time) and technical noise at each timepoint (which is independent at each timepoint). If there are enough read counts that the central limit theorem applies, we can add the variances:

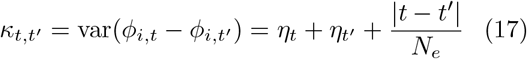

We can measure *κ* for each time interval, and treat the *η*_*t*_ and *N*_*e*_ as unknown parameters that we can fit by minimizing the squared errors of the expected relationship. We can also do this with inverse variance weighting by bootstrapping over barcodes when estimating *κ*.

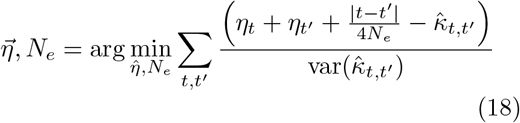

We only use timepoints and barcodes with read count exceeding *r*_*i,t*_ = 50.

We can related these quantities back to the variance in frequency, and obtain an estimate for the dispersion parameter, *c*_*t*_.

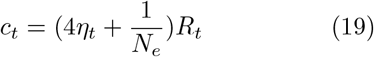

Having obtained our parameter estimates for intermediate parameters, we can finally infer the parameter value for fitness, *s*, using the likelihood function for a negative binomial random variable. The likelihood for a particular value of *s* and *f*_0,*i*_ given a barcode read count of *r*_*i*_, is

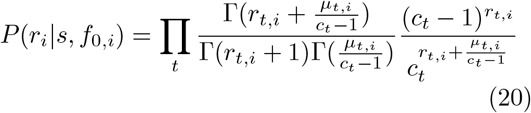

We can marginalize over values of *f*_0_, *i*

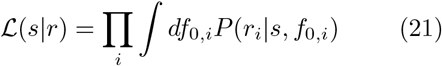

and find the maximum likelihood fitness,

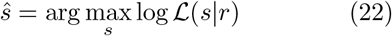

We use a grid search to find the value of *s*. We can use the likelihood function to infer a standard error on our parameter estimation, approximated as the inverse of the square-root Fisher information,

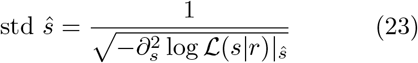

To get a fitness estimate for a single environment, we infer fitness separately for each replicate, and then take a weighted average across our two biological replicates, weighted by the inverse of the inferred standard error.

#### 2. Fitness processing

There were some technical problems in sequencing, resulting in several environments with poor data quality. We dropped these from further analysis. These included all temperature perturbations to the 2 Day base (all Batch 1 conditions), the Batch 4 Salt Base, and 10 *µ*M H89 on the Salt base.

#### 3. PCA on environment space

To assess environmental similarity, we did principal component analysis on our fitness data. First, we took fitness data for all environments and centered and scaled it using `sklearn.preprocessing.StandardScaler`. We then used `sklearn.decomposition.PCA` on the transpose of our standard fitness matrix (which normally has columns as environments and rows as mutants) so that we could understand environmental variation. We projected each environment onto the first two PCs and used `scipy.spatial.ConvexHull` to generate the convex hulls around each base environment.

#### 4. Calculation of z-score

To quantify the magnitude of our perturbations, we used batch replicates as a baseline for variation. Each batch of our fitness measurements included one “unperturbed” base. For each base *B*, we took the collection of batch replicates and calculated the mean fitness for each mutant *i, µ*_*i,B*_. We obtained an observed standard deviation (*σ*_*i,o*_)from these replicates, and we used the average inferred standard deviation from the fitness inference for the replicates 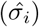, together to represent variation across batch replicates. For each mutant *i* in each environment *e* belonging to base *B*, we calculated a z-score as

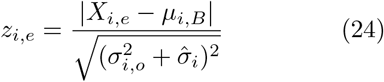

To characterize the entire environment, we took the average *z*_*i,e*_ across all mutants, and the standard deviation across all mutants. For the external bases, we used the same equation for *z*_*i,e*_ as above, but inserted environments corresponding to different bases.

#### 5. Generating the *δX* matrix

To generate the *δX* matrix, we use the numerator of Eq.(24), without taking the absolute value. Thus, for each perturbation *p*, we define

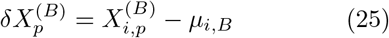

We propagate the error in each measurement (both in generating the standard error of the base environment mean, and in the subtraction of fitness values to generate *δX*_*p*_.

#### 6. Calculating deviation from 1:1 line of perturbations on different base environments

For each mutant, we compared the change in fitness due to the same perturbation, on different base environments (as shown in Fig. 3b). We calculated a deviation from the 1:1 line by subtracting one base’s *δX*_*p*_ from another. We took 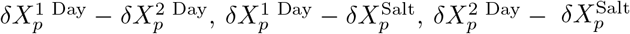, and 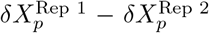. This last comparison was to have a point of reference for the deviation to be expected under biological and measurement noise, and not due to ExE interactions. For each base-base comparison, we only included replicate-replicate *δX*_*p*_ values belonging to one of the two bases in question, to prevent spurious inference of signal from noise.

#### 7. Quantifying deviations from 1:1 line

We used the two sample Kolmogorov-Smirnov statistical test to determine whether the distribution of deviations from the 1:1 line was significantly different between the pairwise base comparisons and the replicate-replicate comparisons. We used `scipy.stats.ks 2samp` to obtain the value for the statistic, and a p-value to assess significance. Additionally, we used the 5th and 95th quantiles to determine how many base-base comparisons were significantly different from the distribution of replicate-replicate comparisons, with a 10% false discovery rate (FDR).

#### 8. Singular value decomposition on *δX* matrices

To generate our model, we use singular value decomposition to factorize the *δX* matrix. For each base, we perform SVD using `numpy.linalg.svd`. We determine fraction of variance explained by component in Fig. 4 by taking each singular value squared, and dividing by the sum of squared singular values.

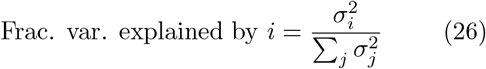

- First, for focal scree plot, we take only the set of adaptive mutants from grant’s paper. Then, we set a standard error threshold (here, we used 0.3), where for a particular mutant, if it has an inferred error on its fitness greater than the threshold, we throw it out. This is to avoid noise causing a huge spike in signal that will corrupt the dimensionality plots. This is of course subject to our choice of threshold, but for a variety of thresholds we find similar numbers. In this case, we throw away 29 mutants out of 258.
- We next take each set of environments, remove No Shake from Salt because we saw that it was not subtle, and do SVD
- We get singular values and calculate fraction of variance explained for each singular value.

#### 9. Calculating a limit of detection for dimensionality

To calculate the number of dimensions that exceed noise in our data, take the same approach as (18). We use the inferred standard error in our fitness estimate, and generate 100 random matrices of the same shape as our data, centered at 0. Each entry is drawn from a normal distribution with mean 0 and standard deviation equal to the inferred standard error at the corresponding index in our data. From this, we can ask how much variation in the true data would have been explained by components derived from the noise-only matrix, by taking the singular values squared from the noise matrix and dividing by the sum of squared singular values in the data. The maximum amount of variation explained by noise matrices represents our limit of detection.

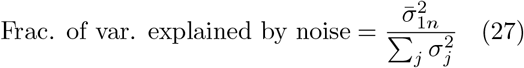

where 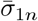 is the average first singular value from each fold of the noise-only matrices, and the *σ*_*j*_ in the denominator come from the *δX* matrix.

#### 10. Linear regression on fitnotypes

On a methodological note, it might seem unnecessary to approach overlap between vectors via prediction, rather than directly looking at some quantity like the inner product, or cosine of the angle between the two 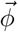 vectors. We take this approach because the method we use to infer our 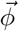 vectors is matrix factorization, which is underdetermined. This means that any rotation or linear transformation of our latent fitnotype space will be equally good at reconstructing our data, so we must focus on properties of our model that are invariant under linear transformation. Prediction error is one such quantity.

We use prediction error for a linear model trained in one base, and tested in held out environments, to asses model success and overlap in latent space. First, we do SVD on *δX* all environments in one base, but we leave out a single perturbation. Let B1 be our focal base, and B2 is a target base. For a perturbation p, we have the reduced matrix 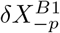, where we have removed perturbation p. We then do SVD on this reduced matrix.

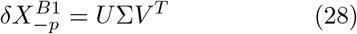

For each value of k up to K=6, we use the k’th column of *U* as a feature in a linear regression, fitting coefficients for the held out perturbation *p* within base, but also for all environments in the target base, *B*2. We can sum the variance from each component, plus the residuals, to equal the total variance, because our regressors are orthogonal (they are singular vectors, after all). This allows us to neatly partition the variance by component. For each value of k, we hold out each perturbation, fit, and generate a prediction for the environments within the focal base, *B*1. The we use the normal *δX*^*B*1^ matrix as the design matrix (without reducing it) for a regression in the target base. We can also ask how much variation is explained within a base if we use SVD with an 8 dimensional model, which is the lowest-error 8 dimensional linear model (11). This represents an ideal, within-training-set error. We can then compare predictive power across components and across bases, as in Fig. 5b and c. Note, within base, we are actually training on slightly different models for each held out prediction, and aggregating predictive power across. By comparing the unexplained variation across bases, and the weights of each component, we can glean information about the underlying fitnotype spaces, even though they are latent.

## DATA AND CODE AVAILABILITY

Source code for the data analysis and figure generation is available at Github (https://github.com/omghosh/limiting-functions). The software repository for the barcode counting code can be found at Venkataram, 2020 (https://github.com/sandeepvenkataram/BarcodeCounter2).

## ACKNOWLEDGMENTS

The authors would like to thank members of the Petrov lab and Good lab for helpful discussions and feedback. We would like to thank Mihkail Tikhonov, Gabriel Mel de Fontenay, Anthony Thomas, Matthew Aguirre, Jonathan Pritchard, Seppe Kuehn, Douglas Stanford, Milo Johnson, Daniel Wong, and Sasha Khristich for helpful discussions. We would like to thank Mikhail Tikhonov, Clare Abreu, Shaili Mathur, Sasha Khristich, Anastasia Lyulina, Vivian Chen, José Aguilar-Rodriguez, Sophie Walton, Manuel Razo, and Zhiru Liu for their help with the experiments (Big Batch Bootcamp team). We would like to thank Rike Stelkens, Kerry Geiler-Samerotte, Hunter Fraser, and Mark Siegal, and Caravaggio Caniglia for feedback on the manuscript. We thank the Stanford Research Computing Center for use of computational resources on the Sherlock cluster. B.H.G. acknowledges support from NIH NIGMS Grant No. R35GM146949. D.A.P. acknowledges support from NIH NIGMS Grant No. 5R35GM118165-07. O.M.G. acknowledges support from the NSF-GRFP. B.H.G. and D.A.P. are Chan Zuckerberg Biohub - San Francisco Investigators.

## SUPPLEMENTAL INFORMATION (SI)

### 1. GENERATING *δX* CORRELATION MATRICES AND ASSESSING EXE INTERACTIONS

To further explore the gxExE interactions we discovered in Section I.B, we asked whether the *δX* of all mutants in two different environments were correlated with each other, and generated a correlation coefficient, or Pearson’s r, with `scipy.stats.pearsonr`. Then, we can specifically zoom in on pairwise comparisons between the same perturbation on different bases, and ask whether these look more or less correlated than different perturbations on the same base. For example, in Fig. S2a, we show the effect of adding the 0.5% ethanol perturbation on top of each base, for all mutants. We find that in general, there is a positive correlation across the different bases. In contrast, when we added 4 uM H89, a drug, to our bases, we find that its effect is actually anti-correlated between the Salt base and the other two bases (Fig. S2b). To capture all of these effects, in Fig. S2c and d, we show the correlation matrices of environment to environment *δX* correlation for all pairs of environments. We first clustered rows of this matrix by perturbation type, and then by batch (panel c). Below, we present the same matrix with a different clustering. We first clustered mutants by base, and then by batch (d). We included these partitions in Fig. S2c and d as dashed lines on top of the correlation matrix. We find that the base-clustered matrix appears to have much more structure, and in particular block diagonal structure, than the perturbation-clustered matrix. This further supports the idea that the effect of environmental perturbations is heavily determined by the base.

To quantify the block-diagonality of each matrix, we used a tool from network theory called the “modularity” score, *Q*. Typically used to quantify the modularity of a network, *Q* can intuitively be understood as a measure of how much more density falls within a partition than would be expected of a random matrix, where the row sums are conserved. First, we scaled the values of the correlation matrices, which initially lay between −1 and 1, to fall between 0 and 1. Then, for each partition of the data, we calculated the difference between the observed and expected weight under a model of randomly distributed weights. As the matrix approaches perfect block-diagonal form, *Q* approaches 1. We calculated *Q* for the scaled correlation matrix with perturbation partitions and base-environment partitions, and found that as expected, *Q* is significantly higher for the base partition. We obtained values of Q=0.35 for the base-clustered matrix, and Q=0.08 for the perturbation clustered matrix.

### 2. SVD ON ADDITIONAL MUTANT SETS

We quantified the dimensionality of the rest of our “biological replicates” to include in Fig. 4c from scree plots like in Fig. 4a. We obtained estimates for detection limit as we described in the main text, using noise-only matrices. In Fig. S3, we present the scree plots that were the source of these estimates.

### 3. ALTERNATIVE WAYS TO QUANTIFY DIMENSIONALITY

We explored two other approaches to quantifying dimensionality. First, instead of allowing each base to have its own limit of detection, we set an overall limit of detection at the maximum fraction of variance explained by any of the noise matrices, yielding a more conservative estimate. In Fig. S4a, we show the inferred dimensionality using this “overall” detection limit. Our dynamic range is much smaller, but the conclusion from the main text does not change. We do not see lower dimensionality in the evolution base.

The number of dimensions that fall above a threshold is one way to quantify “dimensionality,” but it is a discrete value. Another way to capture the scree plots in Fig. S3 is to calculate the entropy of the scree plot for the “real” components. To calculate Shannon entropy, we use the standard formula for the entropy of a distribution,

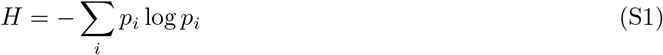

Here, we use the “distribution” of variance explained, so

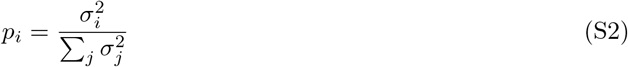

We use only components that fall above the noise limit for the calculation of entropy. Again, we can use either their individual limits of detection, as in the main text, or an overall limit of detection. In Fig. S4b, we use the overall detection limit, and in Fig. S4c, we use the individual detection limit. Using entropy instead of a discrete number does not change the result in the main text, which is that the pleiotropic expansion hypothesis is not supported.

### 4. HOW DOES DIMENSIONALITY DEPEND ON NUMBER OF PERTURBATIONS?

The inferred dimensionality of a matrix via a low-rank approximation clearly depends on the rank of the input matrix. In particular, we have roughly 20 perturbations for each base environment, so we are doing a low-rank approximation of a matrix with at most rank 20. This means that we cannot infer a dimensionality that is higher than the rank. In other words,

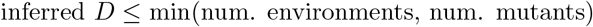

Here, the number of environments is always lower than the number of mutants. To understand the dependence of our dimensionality results in section I.D on the number of environments we included in the study, we randomly selected *n* environments from each base and performed the same procedure described in section I.D. In short, we took 100 random choices of *n* environments for each base, where *n* = 1, …, 16, and did SVD on the sub-matrix for each base. Then we used the inferred noise to generate noise-only matrices, and did SVD on these noise-only matrices. Finally, we asked what fraction the variance in the data could be explained by the top noise component. This sets our limit of detection, and allows us to infer dimensionality. For each instance, we obtain a dimensionality as a function of *n*. In Fig. S5, we plot this inferred dimensionality as a function of *n*, where each point is colored by the base environment.

Often, the dimensionality saturates as we add more environments, supporting the hypothesis that the underlying spaces are indeed low-dimensional. Additionally, across a range of different mutant sets and evolution environments, for any given *n*, we still do not find that the evolution environment is consistently lower dimensional than the other two base environments.

### 5. SYNTHETIC DATA FOR DIFFERENT FITNOTYPE OVERLAP SCENARIOS

To verify our intuition and validate our analyses, we constructed a synthetic dataset assuming that hubs have different levels of overlap in their fitnotypes. We assume that mutations have random effects on a set of *K* latent fitnotypes, drawn from a standard Normal distribution. Each of these *K* fitnotypes will have a random, non-negative weight in an environment, drawn from a uniform distribution between [0, 1). A base environment is characterized by which fitnotypes are relevant, and environmental perturbations will simply perturb the weights of these fitnotypes, but won’t change their identity. In Fig. S6a, we show three example environment-to-fitnotype matrices, corresponding to the different scenarios of fitnotype overlap referenced in Fig. 5. In the first scenario, fitnotypes are equally important in both base environments. In the third scenario (orthogonal fitnotypes), fitnotypes that matter in Base 1 have 0 weight in Base 2, and vice versa. In the partially overlapping fitnotypes scenario, some fitnotypes are shared and others are private to one base or the other. The resulting fitness matrices are shown in Fig. S6b, obtained by multiplying the mutant-to-fitnotype matrix and the environment-to-fitnotype matrix together, and adding normally distributed noise with mean 0 and standard deviation 0.05. The separation between Base 1 and Base 2 grows increasingly obvious as there is less overlap in underlying fitnotypes. In Fig. S6c, we do SVD on each base separately and quantify the variance explained by each component (as we did in Fig. 4a. We can see that the number of relevant fitnotypes in each base is recovered by the elbow technique (finding the kink in the plot), likely due to the true linearity of the underlying data, and the low levels of noise (as opposed to the real data). Finally, we used the same linear regression technique described in Fig. 5a to assess fitnotypic overlap. We can see that qualitatively, we recover what we expect: equivalent prediction for fully overlapping fitnotype spaces, less predictive power for partially overlapping fitnotypes, and little to no prediction for orthogonal fitnotypes.

### 6. FITNOTYPE OVERLAP FOR OTHER MUTANT SETS

In Fig. S7, we perform the same analysis as in Fig. 5 in the main text, but for different groups of mutants. We find different patterns of overlap for different mutants, emphasizing the dependence of our approach on the constituent genotypes. Intriguingly, there are multiple instances where a target base is more predictable than the perturbations within the training base for certain groups of mutants. A further investigation of the genotype dependence of these patterns of overlaps would be interesting, but is outside the scope of this work.

### 7. PREDICTIONS SEPARATED OUT BY PERTURBATION

The linear regression procedure presented in Fig. 5a makes a prediction for *δX* of all mutants in our dataset, in a particular perturbation. That perturbation is either a test perturbation held out from the training base, or a test perturbation from a different base. The overall “predictive power” of our model is aggregated over predictions for all perturbations. In Fig. S8, we show the fraction of variance explained by each component of our model for each test perturbation. We see that indeed there is much heterogeneity in which perturbations are predictable. This suggests that different perturbations have different projections onto the relevant fitnotypes, and therefore are not all probing the same spaces.

### 8. COMPARISONS WITH DIFFERENT MODELS

In Fig. S9a, we compare the prediction accuracy of three different models for *δX* in two example environments. If the pleiotropic shift model is correct, we should be able to use it to make better predictions for *δX* across environmental bases than a model that does not consider fitnotypic diversity, or a model that does not incorporate base-dependence of these fitnotypes. The model that ignores fitnotypes, but has full genotypic and environmental information, could easily be very powerful, because it has information about how genes perform in the test environment. For each mutant included in Fig. 4, we know what the putative adaptive mutation was, and in which gene it landed. In many cases, we also have multiple mutations in the same gene, allowing use to calculate a mean *δX* effect across them. In Fig. S9a, the first column shows the predicted *δX* using the mean across the gene class (see color of point for gene information), plotted against the measured *δX*. The gene model is discrete, and assumes that all mutations in the same gene have the same fitness effect. While the model has a non-zero coefficient of determination, it is not highly predictive, and fails to resolve fitness differences between mutations that land in the same gene. The second column of Fig. S9a uses the *δX* effect for the perturbation in question, averaged across the other two bases. We saw in Fig. 3 that using *δX* from one base to predict *δX* in another base was not very accurate, but perhaps smoothing out the heterogeneity from different bases could make this a better predictor? This model is less discrete and can resolve different *δX* effects from mutations in the same gene, but it is even less predictive than the gene model overall.

Both models, however, do much worse at predicting *δX* in these two environments than a full linear model that allows for phenotypic diversity, and allows for re-weighting these phenotypes. In the final column, we use a model very similar to the one described in Eq. 5, but we actually predict the *δX* effect of held-out mutants using a bi-cross-validation approach (18, 36). Operationally, this means that the model is trained on neither the exact environment nor the exact mutants for whom we are attempting to predict *δX*. When we compare these models across all environments (Fig. 5d, we find that there is heterogeneity in the coefficient of determination, but on average our bi-cross-validation model does best at predicting *δX*. It is both more able to resolve subtle differences in fitness effect from mutations in the same gene, and it is able to flexibly re-weight the importance of fitnotypes across environments.

**FIG. S1:**
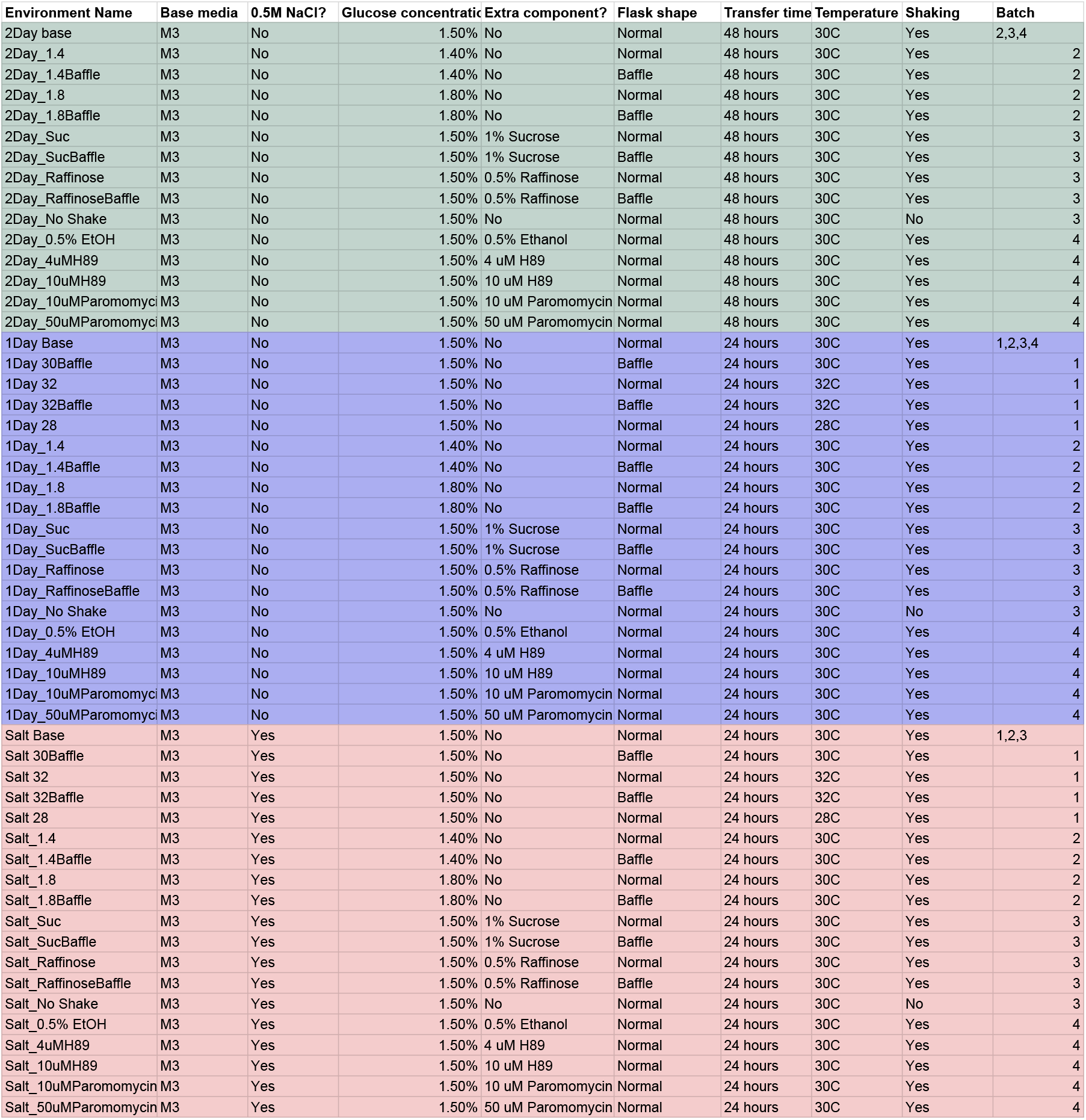
Table of environmental perturbation components. Here, we present a detailed description of each environment and perturbation. We color the rows by base environment, and specify all variants of the environment, along with which batch of the experiment included each perturbation.

**FIG. S2:**
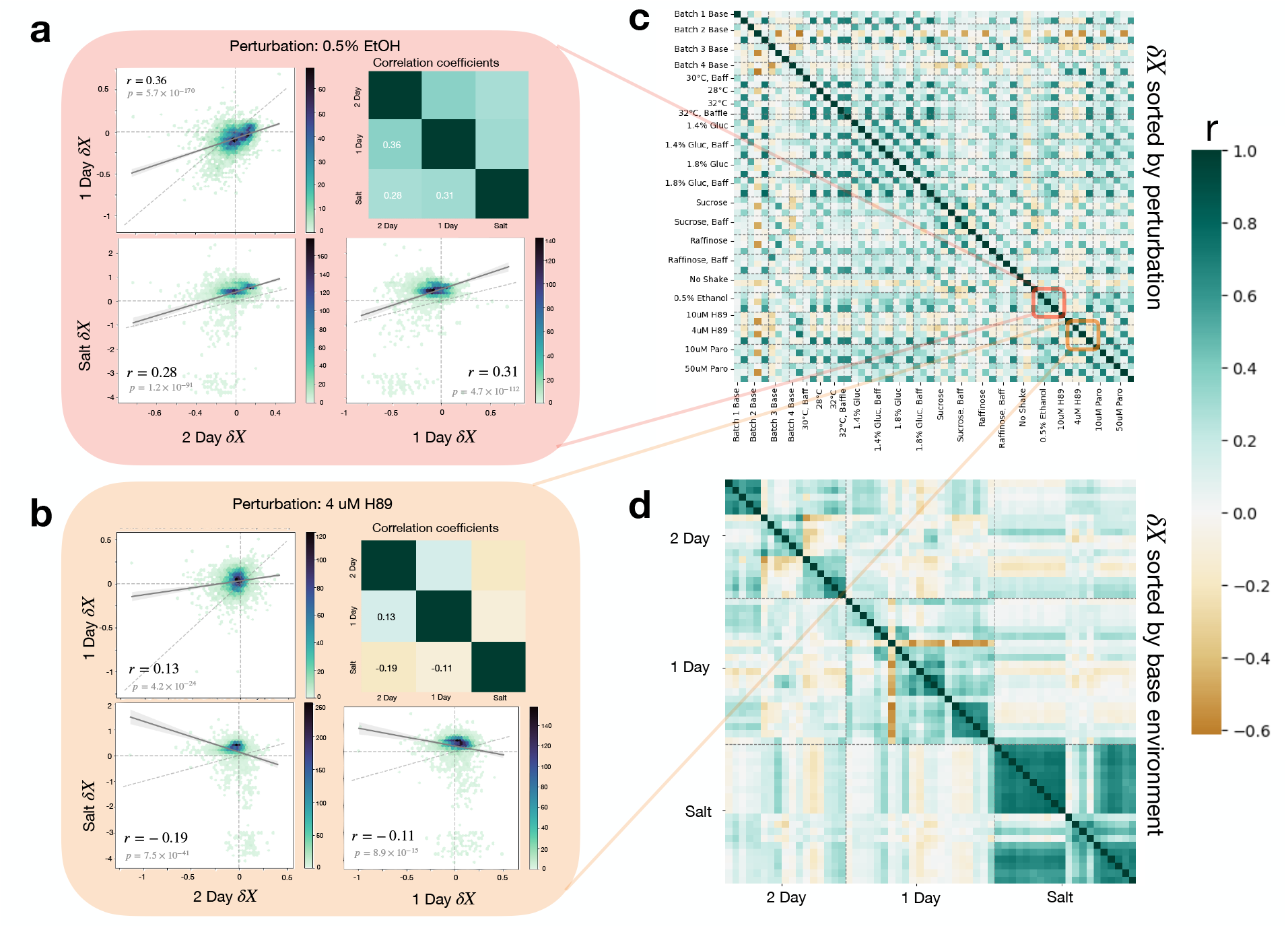
ExE interactions across all mutants. (a) 2 dimensional histograms of *δX* due to focal perturbation on two different base environments. Color of pixel corresponds to number of mutants in the bin. Top right shows correlation coefficient for each environmental comparison (b) Correlation matrix between environmental perturbations, clustered by perturbation (then batch) (top) and base environment (bottom). Block diagonal form is more apparent on the bottom, suggesting that the base environment is important for determining *δX*.

**FIG. S3:**
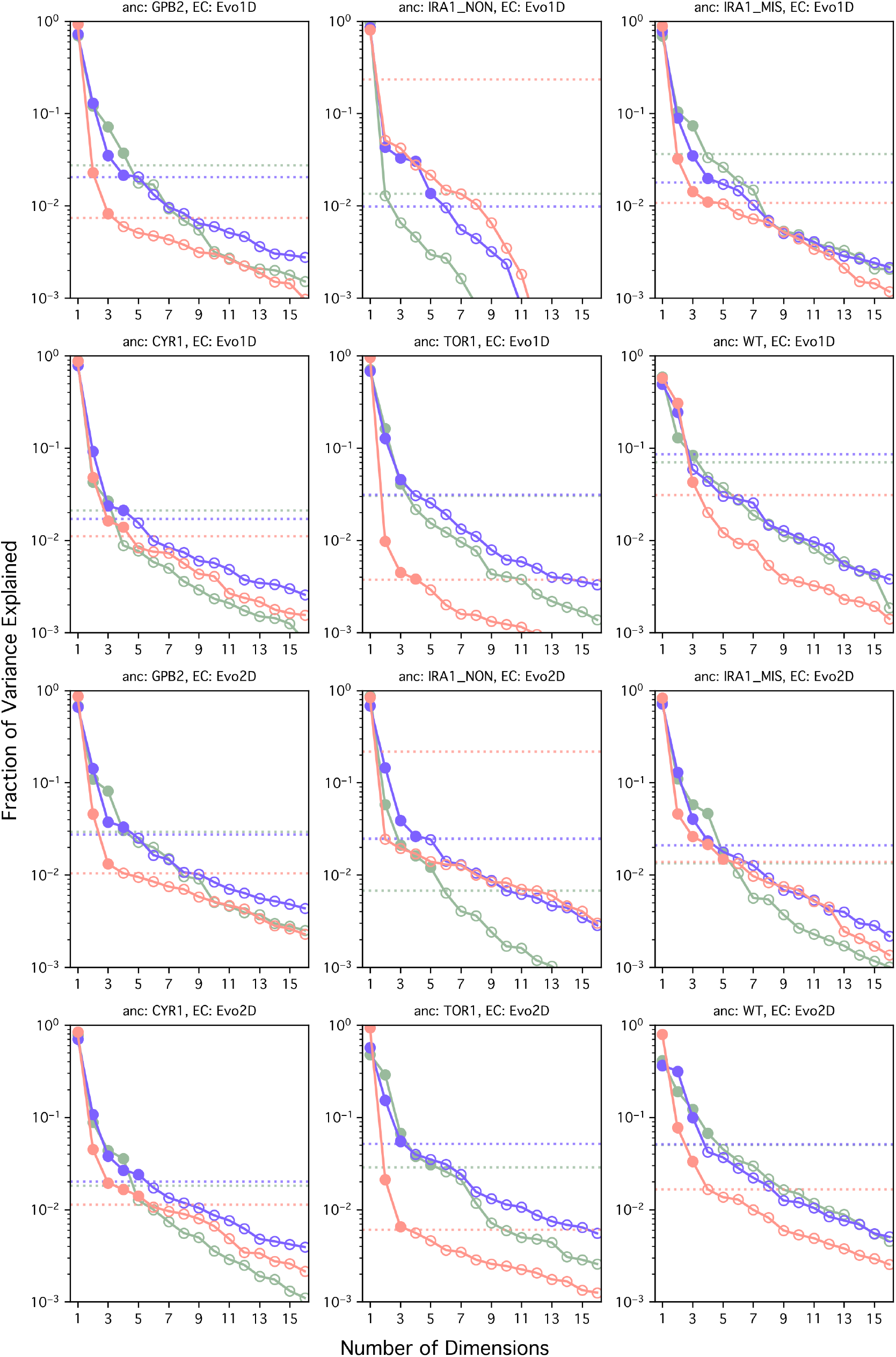
Scree plots for second step mutants. Each set of mutants has a different ancestor, and a different evolution condition. We did SVD on each set of mutants in each base environment, and here show the fraction of variance explained by each component for each base. Green is 2 Day, blue is 1 Day, and pink is Salt.

**FIG. S4:**
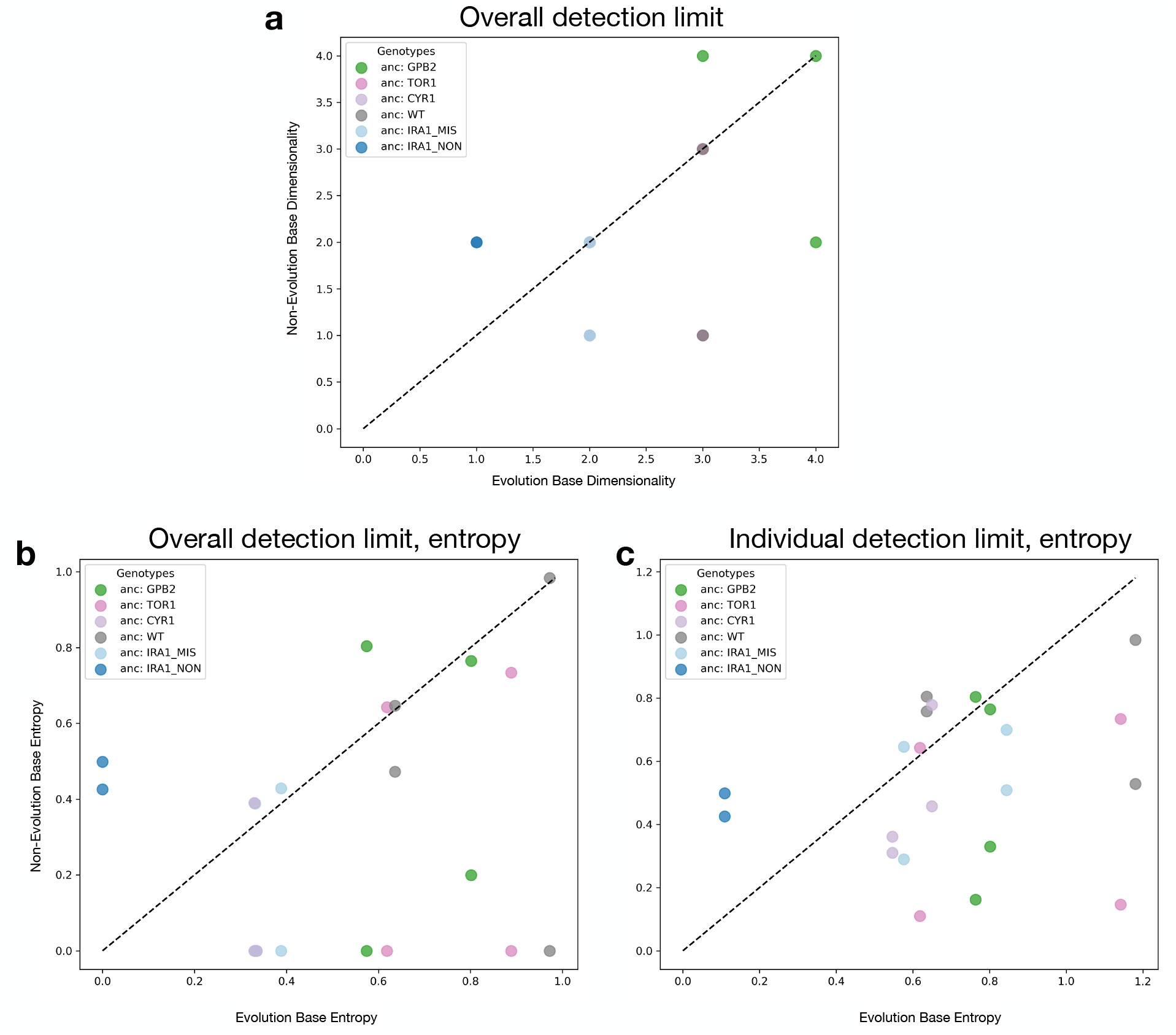
Quantifying dimensionality in alternate ways. Inferred dimensionality in evolution environment base plotted versus dimensionality inferred in alternate base. (a) Dimensionality is inferred based on how many components fall above the most explanatory noise-only matrix for all the bases. (b) Entropy of the distribution of variance explained for the components that fall above the overall limit of detection across bases is used as a proxy for dimensionality. (c) Entropy of distribution of variance explained for components that explain more than each base’s individual detection limit is used as a proxy for dimensionality.

**FIG. S5:**
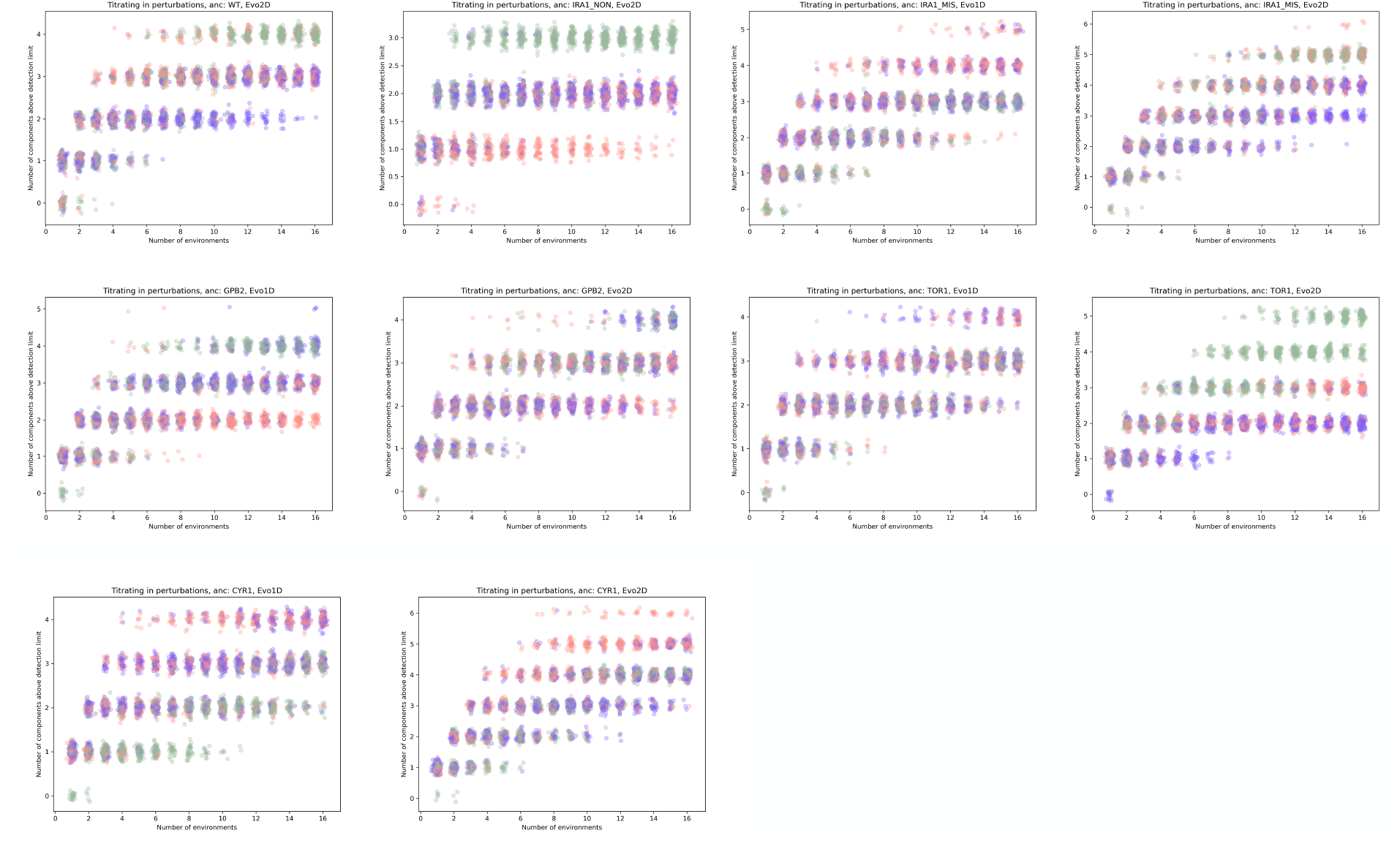
Titrating in perturbations and inferring dimensionality. Each panel is a distinct set of mutants, evolved from a different ancestor and in a different environment. Across a range of n, we still find that the evolution condition is not systematically lower in inferred dimensionality than the other two base environments. X and Y jitter added.

**FIG. S6:**
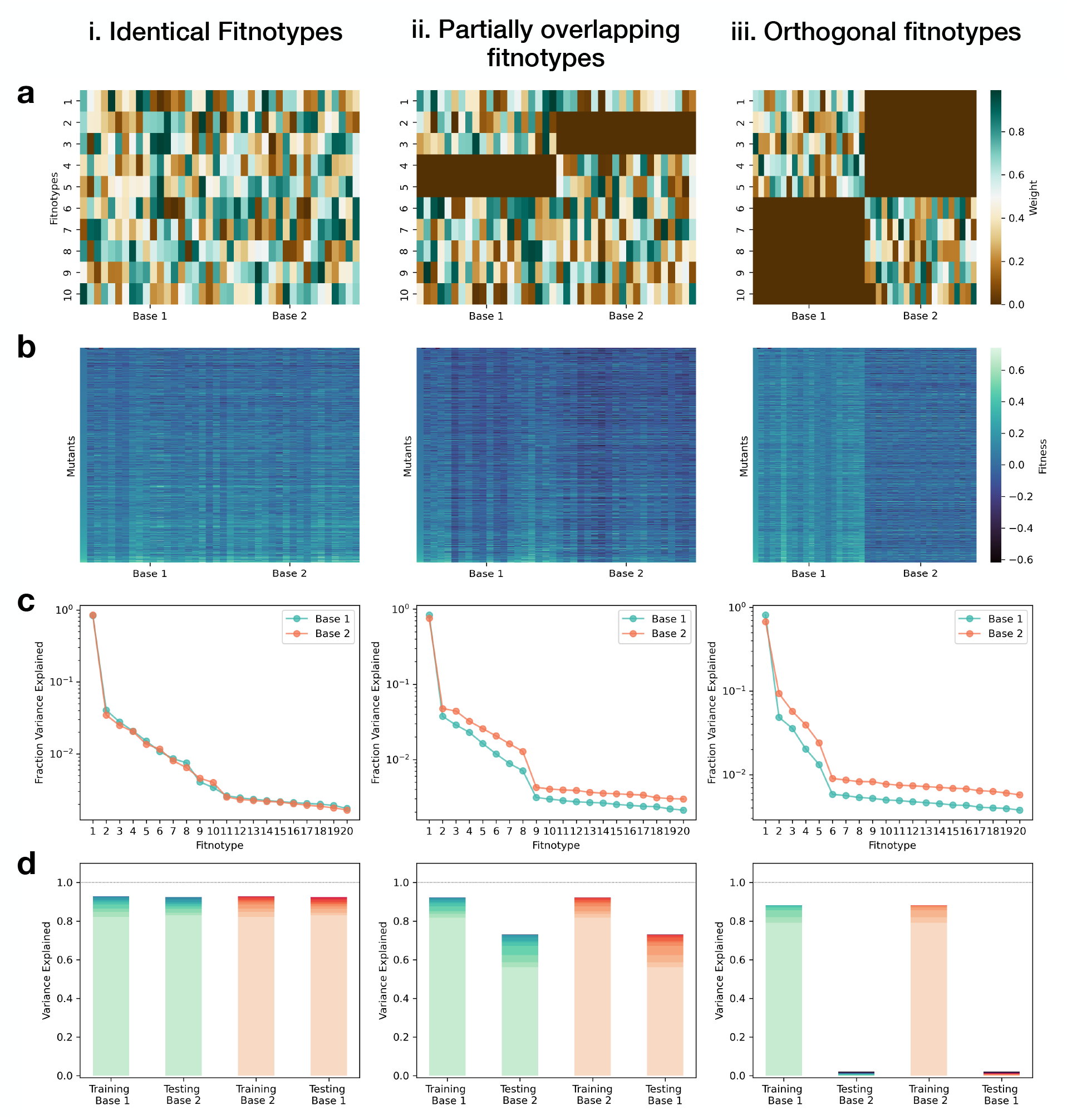
Synthetic data. (a) Underlying weights of fitnotypes (rows) in different environments (columns) for different scenarios of fitnotype overlap. (b) Fitness matrices for the same mutant-to-fitntotype matrix, but different environment-to-fitnotype matrices (from panel a). (c) Variance explained by each inferred fitnotype using SVD to identify fitnotypes. (d) Prediction within and across bases for different fitntoype overlap.

**FIG. S7:**
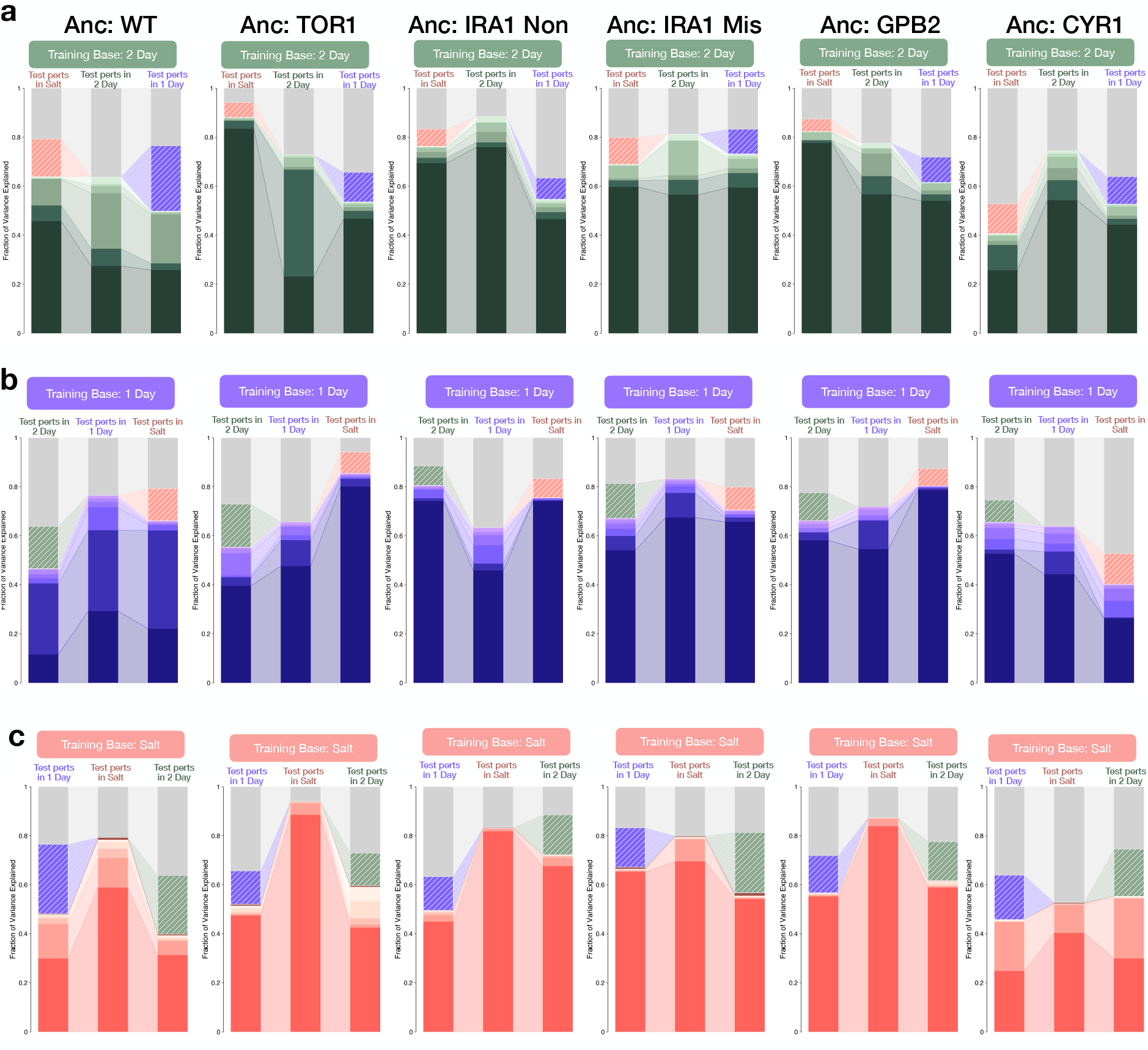
Predicting *δX* with linear regression for different mutant sets. Columns correspond to predictions for different mutant sets (mutants that evolved from different ancestors). We show results for training base 2 Day (a), training base 1 Day (b), and training base Salt (c).

**FIG. S8:**
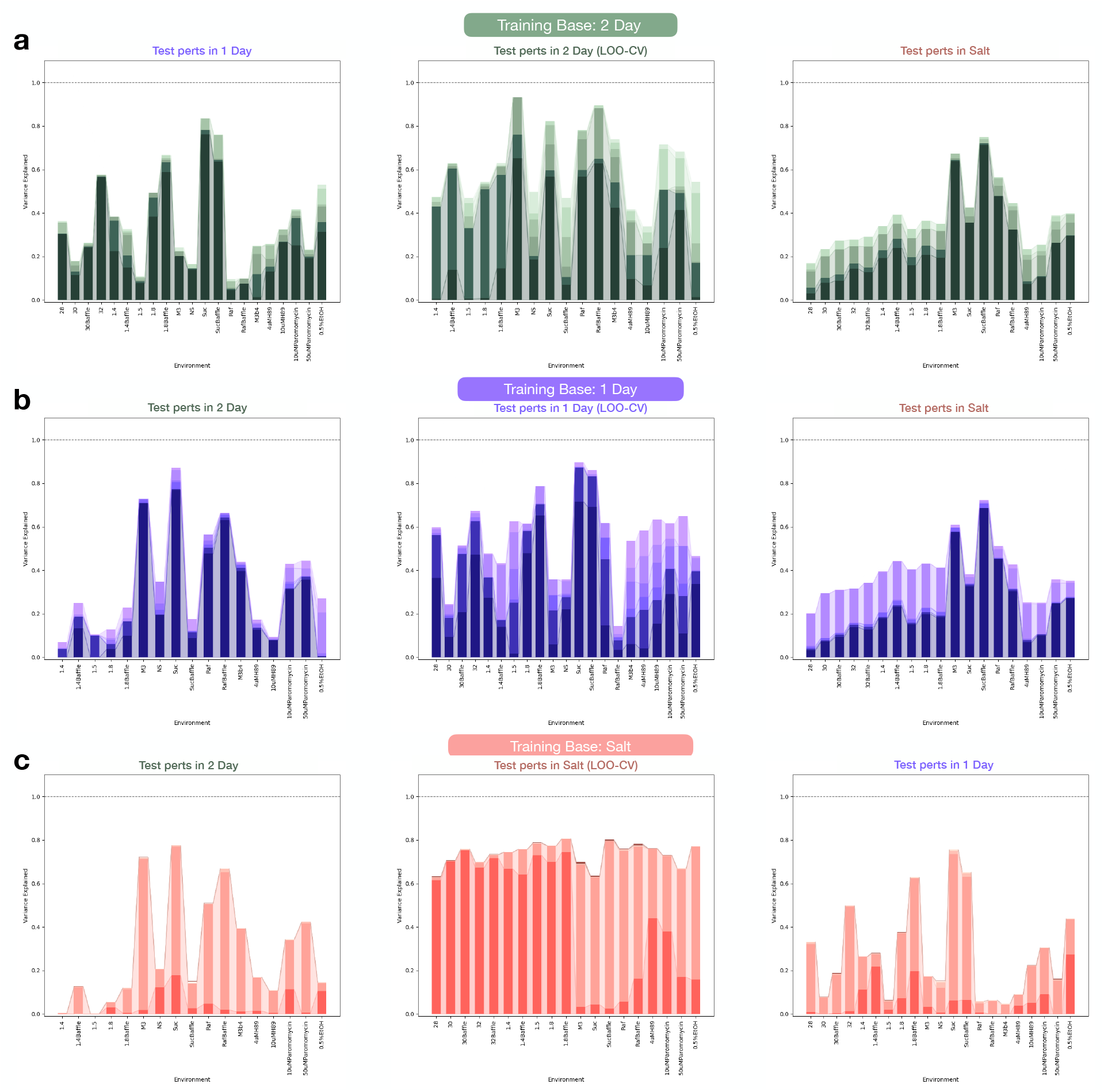
Predictions for target bases separated out by perturbation. Each prediction is done for one perturbation. The contribution of each component from the training base to predicting *δX* in the test perturbation is shown here. (a) Training base is 2 Day. (b) Training base is 1 Day. (c) Training base is Salt.

**FIG. S9:**
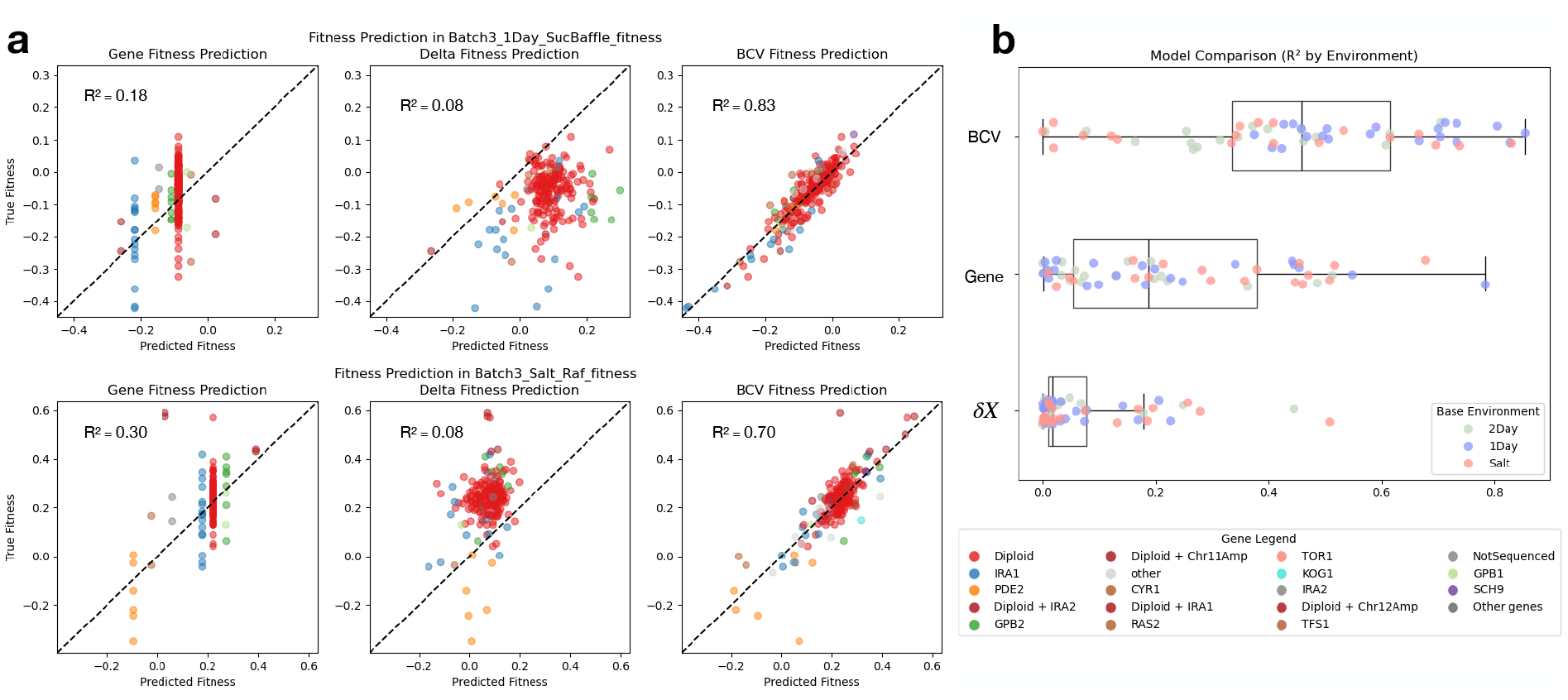
(a) Comparison of model predictions and measured *δX* for 2 Day adaptive mutants, using a gene-only model, a perturbation-only model, and a linear fitnotype model (BCV), for two environments. (b) R^2^ for all environments for each model, colored by base environment. The *δX* is least predictive on average, and the BCV is most predictive on average, despite heterogeneity.

